# A single-cell transcriptome atlas of the adult muscle precursors uncovers early events in fiber-type divergence in *Drosophila*

**DOI:** 10.1101/806281

**Authors:** Maria Paula Zappia, Lucia de Castro, Majd M. Ariss, Abul B.M.M.K. Islam, Maxim V Frolov

## Abstract

In *Drosophila*, the wing disc-associated adult muscle precursors (AMPs) give rise to the fibrillar indirect flight muscles (IFM) and the tubular direct flight muscles (DFM). To understand early transcriptional events underlying this muscle diversification, we performed single cell RNA-sequencing experiments and built a cell atlas of AMPs associated with third instar larval wing disc. Our analysis identified distinct transcriptional signatures for IFM and DFM precursors that underlie the molecular basis of their divergence. The atlas further revealed various states of differentiation of AMPs, thus illustrating previously unappreciated spatial and temporal heterogeneity among them. We identified and validated novel markers for both IFM and DFM precursors at various states of differentiation by immunofluorescence and genetic tracing experiments. Finally, we performed a systematic genetic screen using a panel of markers from the reference cell atlas as an entry point and found a novel gene, *Ama*, which is functionally important in muscle development. Thus, our work provides a framework of leveraging scRNA-seq for gene discovery and therefore, this strategy can be applied to other scRNA-seq datasets.

## Introduction

Muscle fibers exhibit significant variability in the biochemical, mechanical, and metabolic properties, which are defined by the needs and specialized functions of each muscle. The *Drosophila* adult skeletal muscle represents an ideal system to dissect the transcriptional events regulating muscle diversity. In the adult fly, the thoracic muscle contains two types of flight muscles, the indirect flight muscles (IFM) and the direct flight muscles (DFM), that present distinct structure, positioning, patterning and specialized function (Lawrence, 1982). The IFM are fibrillar muscles that provide power to flight, whereas DFM are tubular muscles required for proper wing positioning. Fiber fate is specified by the transcriptional factors *extradenticle* (*exd*), *homothorax* (*hth*) and *spalt major* (*salm)* which control the expression of fiber-specific structural genes and sarcomeric components during myofibrillogenesis at early pupal stages (Bryantsev et al., 2012; Schönbauer et al., 2011). However, there is very little information about the extent of divergence of the transcription programs in the muscle precursor cells that give rise to these two muscle types.

The adult muscle precursor (AMP) cells that give rise to both IFM and DFM are specified early in development and associated with the wing imaginal discs (Bate et al., 1991; Dobi et al., 2015). The AMPs are considered muscle-committed transient stem cells and share some features with the vertebrate adult muscle stem cells called satellite cells (Figeac et al., 2007). During larval stages, AMPs undergo extensive proliferation to reach a population size of 2,500 cells by the late third instar larval stage (Gunage et al., 2014). Interestingly, the AMP cells that will form the IFM, are located on the presumptive notum and show high levels of expression of both *vestigial* (*vg*) and *cut* (*ct*), whereas the AMPs that will give rise to the DFM are located near the presumptive wing hinge and only show very high levels of expression of *ct* but not *vg* (Sudarsan et al., 2001) (Figure 1A). It has been suggested that such AMP divergence is maintained by both intrinsic and extrinsic signals, the latter emanating from epithelial cells of the wing disc. One such signal is wingless (wg), secreted from the notum, that maintains *vg* expression in IFM precursors and establishes a boundary between IFM and DFM myoblasts (Sudarsan et al., 2001). However, with the exception of *ct* and *vg*, no other genes are known to distinguish these two groups of AMPs, raising the question of whether changes in gene expression are limited to these two genes. Compounding the issue is the lack of knowledge about the level of heterogeneity within each group of AMP cells. Yet, this is important for the interpretation of the experiments in which transplantation of the labeled wing disc-associated AMP cells into larval hosts led to an indiscriminate contribution to the developing adult muscles. It was suggested that the specification of AMPs at larval stage is not definite yet, and therefore, can still adapt to changing environmental cues (Lawrence and Brower, 1982). Whether this conclusion is applicable to an entire pool of AMPs or to a more naïve population of AMPs that is uniquely capable of such transformation is unknown.

**Figure 1.**
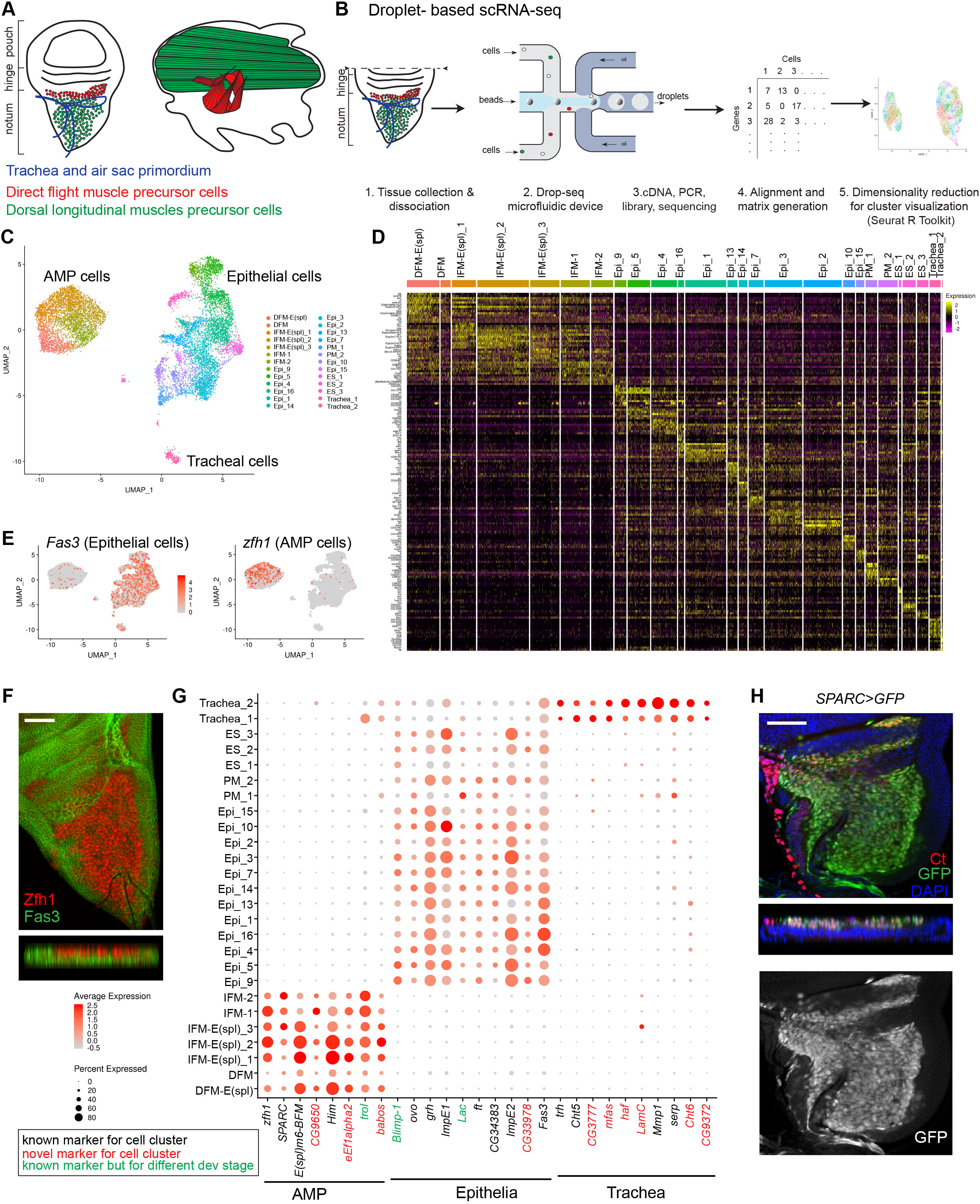
Single-cell atlas of the wing imaginal disc identifies diverse cell types. (A) Diagram of the origin of the flight muscle. Left panel: there are two types of adult muscle precursors (AMPs) that give rise to the flight muscles, the IFM precursors labeled in green and the DFM precursors in red. These are located in the adepithelial layer overlaying the epithelium of the wing disc. In addition to the AMPs and the epithelial cells, both trachea and air sac primordium are in close association with the wing discs, as indicated in blue. Right panel: Lateral view of the adult flight muscles arrangement in the thorax. IFM are in green and DFM in red. (B) Illustration of the experimental workflow for single-cell analyses. Third instar larval wing discs were dissected, pouch was removed, and dissociate into single-cell suspension. Droplets containing unique barcoded-beads and single cells were collected. Following library preparation, sequencing data were aligned, and a gene-cell expression matrix was generated and analyzed using Seurat for identification of variable genes and unsupervised cell clustering based on gene expression similarity. (C) Annotated cell-type, including 6,983 epithelial, 272 tracheal and 4,544 AMP cells, in UMAP plot of the reference single cell atlas of wild type notum and hinge wing discs at third instar larval stage. (D) RNA expression heatmap showing the top differentially expressed gene markers for each cluster of the reference single cell atlas dataset. Cells (column) are clustered by the expression of the main marker genes (row). (E) Average expression level of the genes *Fasciclin 3* (*Fas3*, left panel) and *zfh1* (right panel) used as markers to assign epithelial and AMP cells, respectively, in the reference atlas dataset. (F) Confocal single plane image of third instar larval wing disc and orthogonal views of the disc illustrating the layering and distribution of myoblasts and epithelial cells within the notum, stained with anti-Zinc finger homeodomain 1 (Zfh1, red) to mark the AMP cells and anti-Fasciclin 3 (Fas3, green) to mark the epithelial cells. Scale bars represent 50 μm. (G) Dot plots showing the expression levels of the marker genes identified for the AMP, epithelial and tracheal cells across the 26 clusters of the reference cell atlas. Color intensity represents the average normalized expression level. Dot diameter represents the fraction of cells expressing each gene in each cluster. (H) Confocal single plane image of third instar larval wing disc and orthogonal view of the *SPARC>GFP* (green) notum disc stained with anti-Cut (Ct, red) to mark the AMPs and 4,6-diamidino-2-phenylindole (DAPI, nuclear marker, blue). Scale bars represent 50 μm. Full genotype *y-, w-/w-; UAS-GFP/+; SPARC-GAL4/+*.

To investigate the early divergence in the transcriptional programs between DFM and IFM in proliferating AMPs, we performed single cell RNA-seq experiments and constructed a high-resolution reference cell atlas comprising 4,544 AMP cells, which yield 1.8x cellular coverage. We found that IFM and DFM precursors have distinct transcriptional signatures indicating that the genetic regulatory networks driving each muscle type diverge prior to fiber fate specification. Unexpectedly, the atlas revealed that IFM and DFM precursors are highly heterogeneous and each group contains distinct populations representing cells at various states of differentiation. Finally, by combining the scRNA-seq approach with an RNAi based genetic screen and genetic tracing experiments, we identified new genes that are important for skeletal muscle development.

## Results

### A single-cell atlas of the wing imaginal disc identifies diverse cell types

We performed scRNA-seq to identify the differences in the transcriptional profiles of two subtypes of AMPs that give rise to direct and indirect flight muscles. Third instar larval wild type wing discs were dissected, cut along the presumptive hinge to remove most of the wing pouch and enrich the sample for AMPs that are located in the adepithelial layer of the notum (Figure 1A). Dissected tissue was dissociated into single-cell suspension and processed using the Drop-seq protocol (Macosko et al., 2015a). Single-cell transcriptomes of eight independent replicates of two wild type stocks *1151-GAL4* and *1151>mCherry-RNAi* were sequenced, and data were processed using an integrative analysis in Seurat 3 package (Stuart et al., 2019). After filtering poor quality cells, Uniform Manifold Approximation and Projection (UMAP) dimensionality reduction algorithm was used to visualize cell populations (Figure 1B, see Methods for more details). To eliminate batch effects, clusters of cells that were not evenly represented among the replicates were removed (Figure S1A-B). Using these stringent criteria, we retained 11,527 high quality cells to generate a reference wild type atlas of the wing imaginal disc (Figure S1C).

Unsupervised graph-based clustering identified 24 cell clusters (Figure 1C), each exhibiting a distinct gene expression signature (Figure 1D, Supplemental Table S1, Supplemental Table S2). Nineteen clusters represented epithelial cells based upon the expression of the epithelial marker *Fasciclin 3* (*Fas3)* (Bate and Martinez Arias, 1991) (Figure 1E). The remaining seven clusters comprised 4,544 cells that lacked *Fas3* expression but showed high levels of the myoblast-specific genes: *Zn finger homeodomain 1* (*zfh1*) and *Holes in muscle* (*Him*) (Lai et al., 1991; Soler and Taylor, 2009) indicating that these are AMPs. Accordingly, the principal component analysis revealed that the majority of gene expression variance among the cells was accounted by epithelial-and muscle-specific genes (PC1, Figure S1D). Using anti-Fas3 and anti-Zfh1 antibodies, epithelial cells and AMPs were visualized in wing discs by immunofluorescence (Figure 1F).

To further explore the differences between AMP and epithelial clusters, we examined the list of differentially expressed genes between these two cell types (Supplemental Table S3). In addition to the canonical AMP-specific markers, *Him* and *zfh1*, other muscle-related genes, such as *Secreted protein, acidic, cysteine-rich* (*SPARC*), *CG9650*, *eukaryotic translation elongation factor 1 alpha 2* (*eEf1alpha2*), *terribly reduced optic lobes* (*trol*) and *babos* were found to be highly expressed in the AMP clusters (Figure 1G). Among them, trol was shown to be expressed in muscle attachment sites in embryonic muscle (Friedrich et al., 2000) and *BM-40-SPARC* in the AMP in third instar larvae (Butler et al., 2003). We confirmed this result using the *SPARC>GFP* reporter line (Figure 1H).

The assignment of the epithelial clusters was further supported by the specific expression of *ovo*, *grainy head* (*grh)*, *Ecdysone-inducible gene E1* (*ImpE1)*, *four-jointed* (*ft*), *Ecdysone-inducible gene E2* (*ImpE2*) and *CG34383* (*Kramer*) genes (Figure 1G), which were reported to be expressed in diverse epithelium, and *Blimp-1* and *Lachesin* (*Lac*) in trachea epithelium (Llimargas et al., 2004; Ng et al., 2006). We also confirmed the expression of *grh* in the epithelial layer of the wing disc directly using *grh::GFP* reporter line and showed mutual exclusivity of *grh::GFP* expression with the myoblast marker twist (Supplemental Figure S1E).

Two epithelial clusters (Trachea_1 and Trachea_2) expressed a canonical tracheal marker *trachealess* (*trh*) (Sato and Kornberg, 2002) indicating that these clusters contain tracheal cells. Accordingly, we detected the expression of several known tracheal-specific genes including *Chitinase 5* (*Cht5*), *serpentine* (*serp*) and *Matrix metalloproteinase 1* (*Mmp1*). Interestingly, *pebbled* (*peb*) and *breathless* (*btl*) were expressed exclusively in cells of Trachea_1, while *Ultrabithorax* (*Ubx*) (Brower, 1987), *Gasp*, *Cuticular protein 49Ag* (*Cpr49Ag*), *Pherokine 3* (*Phk-3*), *Cuticular protein 12A* (*Cpr12A*) and *Cystatin-like* (*Cys*) exhibited Trachea_2-specific expression (Figure S1F). We also identified novel markers for each cluster (Figure 1G, Figure S1F, Supplemental Table S4).

We concluded that the single-cell reference atlas contains epithelial cells of the wing disc, adult myoblast precursors and tracheal cells, and therefore, accurately represents the cellular diversity of the wing imaginal disc.

### Cells in the epithelial clusters map to spatially distinct regions of the wing disc

Unbiased clustering analysis grouped epithelial cells of the wing disc into 17 transcriptionally distinct cell clusters. To assign the spatial position of individual clusters in the wing disc, we selected the marker genes for each cluster, and then searched the literature for published *in situ* expression patterns of these genes in the wing disc. The cluster positions were then mapped to the presumptive adult structures using the cell fate map of the wing disc (Bryant, 1975)(Figure 2A). In this way, we assigned the identities of ten clusters to the disc proper, two clusters to the peripodial membrane and three clusters to cells associated with external sensory organs (Figure 2B-C). The corresponding markers used for assignment as well as the new markers are shown in feature map (Figure 2D, Figure S2A, Supplemental Table S2) and in dot plot (Figure 2E).

**Figure 2.**
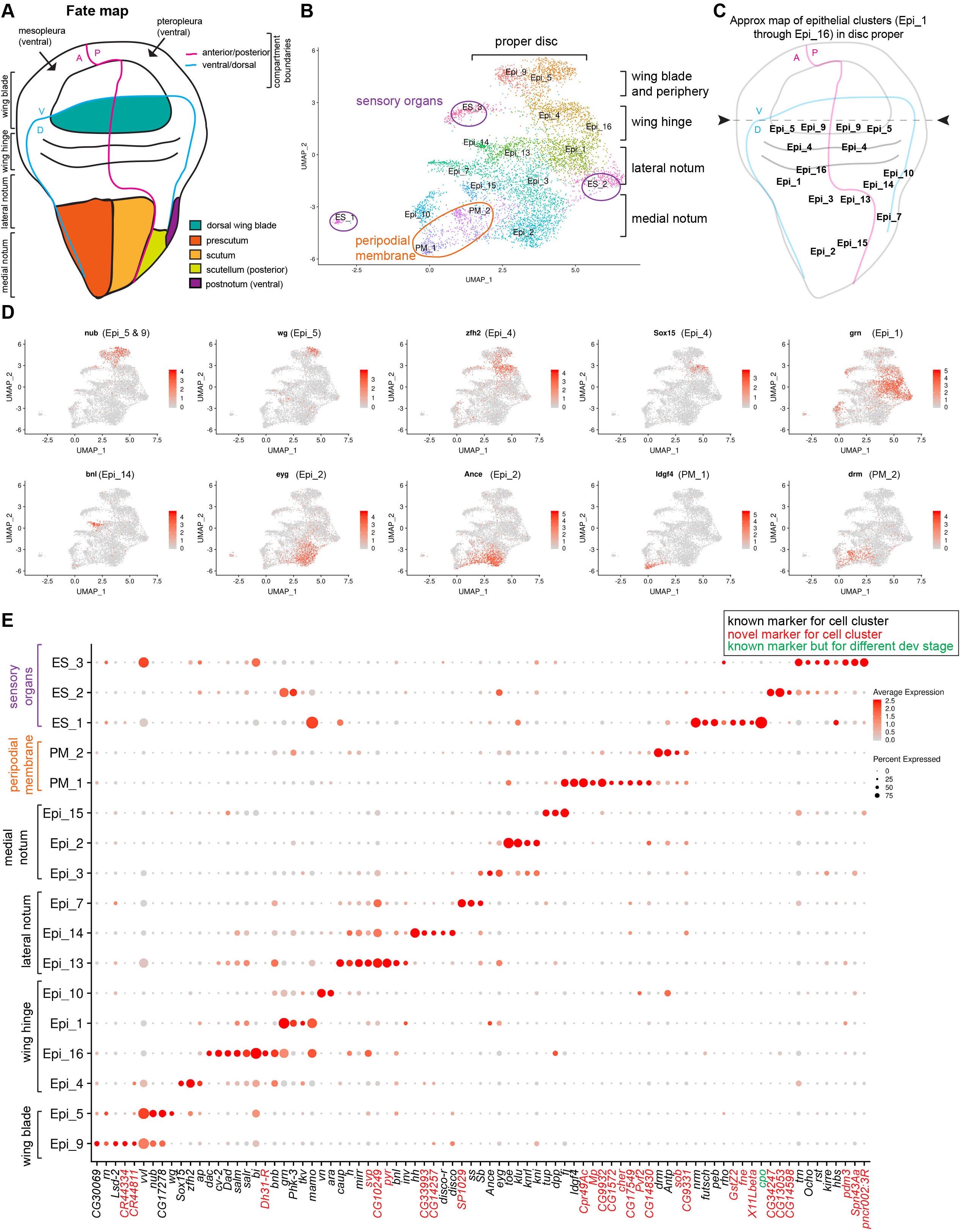
Cells in epithelial clusters map to spatially distinct regions of the wing disc. (A) Schematic representation of the wing imaginal disc and its relationship to the adult wing. The cell fate map of the wing disc is adapted from (Bryant, 1975) (B) Subset of 6,983 epithelial cells in UMAP plot of the wild type reference single cell atlas dataset. (C) Approximate map of the epithelial cluster identified in the reference single cell atlas dataset to distinct position over the disc proper. (D) Average expression level of the genes used as known markers to assign each epithelial cluster in the reference atlas dataset. Cells colored by the expression of the *nub*, *wg*, *zfh2*, *Sox15*, *grn*, *bnl*, *eyg*, *Ance*, *Idg4* and *drm* (E) Dot plots showing the expression levels of the top marker genes identified for the epithelial cells across the 17 clusters of the reference cell atlas dataset. Color intensity represents the average normalized expression level. Dot diameter represents the fraction of cells expressing each gene in each cluster.

Epi_9 and Epi_5 corresponded to the wing blade and the periphery of the wing blade, respectively. This is based on the expression of *nubbin* (*nub*) (Ng et al., 1995), *rotund* (*rn*) (St Pierre et al., 2002), *ventral veins lacking* (*vvl*) (de Celis et al., 1995) and *CG17278* (Mohit et al., 2006) in both clusters, and on the expression of *wingless* (*wg*), which is expressed in the inner ring of the periphery of the wing blade (Couso et al., 1993; Terriente et al., 2008), which was found only in Epi_5. Accordingly, *CG30069* and *Lipid storage droplet-2* (*Lsd-2*), which are expressed in wing blade (Butler et al., 2003; Fauny et al., 2005), were Epi_9 specific markers, while *Zn finger homeodomain 2* (*zfh2*), which is found at the periphery of the wing blade (Whitworth and Russell, 2003), was an Epi_5 specific marker.

Both Epi_4 and Epi_16 mapped to the wing hinge, with Epi_4 located distal to Epi_16. This is due to the expression of *Sox box protein 15* (*Sox15*), the top marker for Epi_4, that is restricted to the hinge (Crémazy et al., 2001) between the inner and outer rings (Dichtel-Danjoy et al., 2009), along with *zfh2* and *apterous* (*ap*) (Cohen et al., 1992). In contrast, cells of Epi_16 expressed high levels of *dachshund* (*dac*) (Mardon et al., 1994), *crossveinless 2* (*cv-2*) (Conley et al., 2000), *Daughters against dpp* (*Dad*) (Tabata et al., 1997), *salm*/*spalt-related* (*salr*) (de Celis et al., 1996a), and *bifid* (*bi*) (Sun et al., 1995). The expression pattern of these genes mostly overlaps in the central area of the hinge along the A/P boundary close to the presumptive lateral notum. Accordingly, a subset of cells of this cluster showed low levels of expression of *araucan* (*ara*) (Gómez-Skarmeta et al., 1996) and *hairy* (*h*) (Usui et al., 2008), which are localized to the lateral notum. Since *decapentaplegic* (*dpp*) was also expressed in Epi_16, and is a marker of the anterior cells in the A/P boundary (Posakony et al., 1991), we reasoned that the cells of cluster Epi_16 are mostly positioned in the anterior compartment near the A/P boundary of the hinge.

Epi_1 was likely localized in the anterior hinge close to the presumptive lateral notum. These cells showed high levels of *grain* (*grn*), which is expressed in the hinge (Brown and Castelli-Gair Hombría, 2000), *thickveins* (*tkv*), which is found in the most anterior and posterior area of presumptive notum (Brummel et al., 1994), and *Phk-3*, which is expressed in the anterior hinge and presumptive notum (Klebes et al., 2005). Epi_10 only expressed high levels of two genes, *knirps* (*kni*) and *knirps-like* (*knrl*), which are found in the posterior area of the hinge (Lunde et al., 1998).

Epi_13 and Epi_14 were localized along the lateral heminotum near the hinge as they share common markers, such as *ara*, *caupolican* (*caup*) (Gómez-Skarmeta et al., 1996), and *mirror* (*mirr*) (Kehl et al., 1998). Epi_13 showed specific expression of *vein* (*vn*) (Simcox et al., 1996) and high levels of *h* (Usui et al., 2008), whereas Epi_14 showed specific expression of the posterior markers *invected* (*inv*) (Coleman et al., 1987) and *hedgehog* (*hh*) (Tabata and Kornberg, 1994), and *branchless* (*bnl*), which straddles the A/P compartment border near the scutum and hinge (Sato and Kornberg, 2002). Thus, the approximate location of Epi_13 was near the presumptive scutum in the lateral heminotum (near scutellum) and Epi_14 was likely in the posterior lateral heminotum (near scutum). Epi_7 approximate location was near the postnotum in the posterior heminotum since the expression of Epi_7 marker *disco-r* is restricted to the ventral edge of the wing (Grubbs et al., 2013) and the expression of another Epi_7 marker *disco* is restricted to the region giving rise to the post-alar bristles near the presumptive lateral scutum (Cohen et al., 1991).

Known markers for the anterior presumptive notum *eyegone* (*eyg*) (Aldaz et al., 2003), *twin of eyg* (*toe*) (Yao et al., 2008), and *klumpfuss* (*klu*) (Klein and Campos-Ortega, 1997) were highly expressed in Epi_2 and Epi_3. Because *ss* (Duncan et al., 1998) and *Stubble* (*Sb*) (Ibrahim et al., 2013) were detected at higher levels in Epi_3, whereas *Angiotensin converting enzyme* (*Ance*) (Siviter et al., 2002) was specifically in Epi_2, we inferred that Epi_3 was positioned in the anterior lateral notum near the hinge and Epi_2 was in the anterior medial notum. The markers for Epi_15 were *four-jointed* (*fj*) and *tailup* (*tup*), which localize to the prospective notum (Cho and Irvine, 2004; de Navascues and Modolell, 2007). Because *dpp*, which was also expressed in Epi_15, downregulates the Iro-C genes (*ara*, *caup* and *mirr*) in the medial notum (Cavodeassi et al., 2002), we reasoned this cluster is most likely positioned anterior to the A/P boundary in the medial notum.

Two clusters, PM_1 and PM_2, corresponded to peripodial membrane, a layer of squamous cells overlaying the epithelial cells of the disc proper. PM_1 was likely mapping to the dorsal peripodial epithelium as it expressed both *Imaginal disc growth factor 4* (*Idgf4*) (Butler et al., 2003) and *Ance*. Two peripodial markers, *drumstick* (*drm*) (Benitez et al., 2009) and *Antennapedia* (*Antp*) (Levine et al., 1983), were highly expressed in PM_2.

The expression of *sca* in ES_1, ES_2 and ES_3 indicated that these clusters represented either developing external sensory organs or neighboring cells contributing to their development (Figure S2B). The expression of *sca* was highest in ES_1, thus indicating that this cluster represents the sensory organ precursor cells (SOP) that are undergoing cytodifferentiation into neuronal receptors. Concordantly, SOP markers *neuromusculin* (*nrm*) (Ghysen and O’Kane, 1989), *futsch* (Klein and Campos-Ortega, 1997), *peb* (Giraldez et al., 2002) and *rhomboid* (*rho*) (Sturtevant et al., 1993) were highly expressed in cells of ES_1. A common marker for ES_2 and ES_3 was *tartan* (*trn*), which is expressed in macrochaete SOP but not in microchaete SOP of the anterior wing margin (Chang et al., 1993), and *Ocho*, which is found in the external sensory organs (Lai et al., 2000). Thus, ES_2 and ES_3 most likely represent bristle precursors cells. However, ES_3 expressed *bifid* (*bi*) and *vvl*, and ES_2 showed high levels of *grn*, *Phk-3* and *Sb*. Therefore, ES_3 was likely surrounding SOP in the presumptive hinge and wing blade and ES_2 in the anterior hinge or anterior presumptive lateral notum. Concordantly, we found *roughest* (*rst*), *kin of irre* (*kirre*) and *hibris* (*hbs*), which are components of the heterophilic Irre Cell Recognition Module associated with cells surrounding SOPs in the anterior wing margin (Linneweber et al., 2015).

Thus, we conclude that unbiased cell clustering performed in Seurat identifies spatially distinct regions of the wing disc.

### The single-cell transcriptome atlas reveals a large compendium of adult muscle precursors

The wing disc-associated adult muscle precursors (AMP) give rise to two distinct sets of flight muscles: indirect flight muscles (IFM) and direct flight muscles (DFM). Therefore, we first assigned the seven AMP clusters by the differential expression of *ct* and *vg* (Figure 3A-B), the only two known markers for DFM and IFM precursors (Sudarsan et al., 2001). Notably, five clusters showed high *vg* and low *ct* pattern of expression, indicating that these represent IFM precursors, while two clusters showed low *vg* and high *ct* expression, and therefore, were classified as DFM precursors (Figure 3B). Seurat analysis further revealed that, similar to *vg*, the level of *zfh1* expression is higher in IFM precursors than in DFM precursors (Figure 3B). Since *zfh1* is considered a general marker for the entire pool of AMP (Lai et al., 1991), we carefully examined zfh1 expression by co-staining the wing disc with anti-Zfh1 antibody and anti-Ct antibody. Figure 3C shows differential expression of Zfh1 and Ct among IFM and DFM precursors, thus, confirming the results of scRNA-seq. Notably, the expression of the flight muscle determinant genes, such as *salm*, *exd* and *hth*, were either too low or unchanged among the AMP clusters.

**Figure 3.**
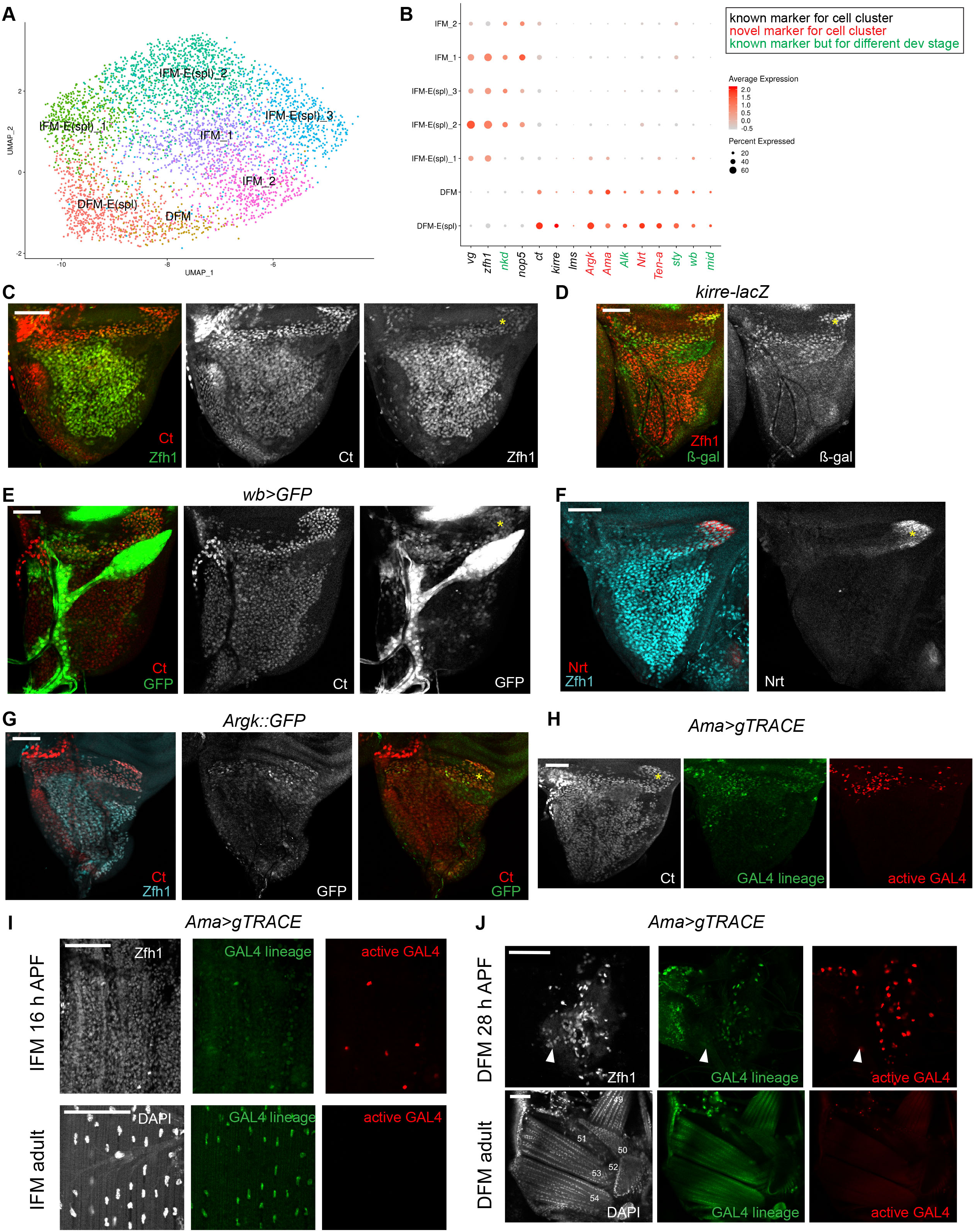
Single-Cell Transcriptome Atlas reveals a large compendium of adult muscle precursors. (A) Subset of 4,544 AMP cells in UMAP plot of the wild type reference single cell atlas dataset. (B) Dot plots showing the expression levels of the top variable genes between IFM and DFM precursors across 7 AMP clusters of the reference cell atlas dataset. Color intensity represents the average normalized expression level. Dot diameter represents the fraction of cells expressing each gene in each cluster. (C, D, E, F, G) Confocal single plane images of third instar larval wing discs (C) stained with anti-Ct (red) and anti-Zfh1 (green), (D) *kirre-lacZ* stained with anti-β-gal (green) and anti-Zfh1 (red), (E) *wb>GFP* (green) stained with anti-Ct (red), (F) stained with anti-Neurotactin (Nrt, red) and anti-Zfh1 (cyan), (G) *Argk::GFP* (green) stained with anti-Ct (red) and anti-Zfh1 (cyan). (H, I, J) Confocal single plane images of different developmental stages and tissues of *Ama>gTRACE* showing the lineage of *Ama-GAL4* (green) and the active GAL4 (red) (H) third instar larval wing disc stained with anti-Ct (white) (I) Forming IFM (specifically, dorsal longitudinal muscle) at 16 h after pupa formation (APF, top panel) and adult (bottom panel) stained with anti-Zfh1 (white) and DAPI (white), respectively. Anterior up. (J) Forming direct flight muscles (DFM) at 28 h APF stained with anti-Zfh1 (white, top panel), and sagittal section of adult DFM stained with DAPI (white, bottom panel), dorsal right. White arrows point to the wing hinge. Anterior up. DFM are numbered in white following (Lawrence, 1982). Yellow asterisks (*) mark DFM area of each notum. Scale bars represent 50 μm. Full genotypes are (C) *1151-GAL4, v-, w-; +; UAS-mCherry-RNAi*, (D) *w-; rp298-lacZ; +*, (E) *w-; wb[MI07688-TG4.1]-GAL4/UAS-GFP; +*, (F) *y-, v-; +; P{CaryP}attP2*, (G) *y-, w-; +; Argk[CB03789]::GFP*, (H,I,J) *w-; UAS-gTRACE/+; Ama[NP1297]-GAL4/+*.

To further examine the differences in the transcriptional programs between DFM and IFM precursors, the expression of the top markers was visualized across cell populations (Figure 3B, Figure S3A, and Supplemental Table S5). There was a large panel of specific markers for DFM precursors. In contrast, only *vg*, *zfh1*, *nop5*, and *naked cuticle* (*nkd*), which is involved in embryonic muscle patterning (Volk and VijayRaghavan, 1994), were highly expressed in IFM-precursor clusters (Figure 3B). Among the DFM-precursor markers, we found *kirre*, a marker of muscle founder cells in DFM precursors (Kozopas and Nusse, 2002). The expression of kirre in DFM precursors, but not in IFM precursors (Figure 3D), is consistent with DFM being formed *de novo*, as opposed to IFM, which lack the specification of founder cells and use larval oblique muscles as templates. We also found *lateral muscles scarcer* (*lms*) expression restricted to DFM precursors, as previously reported (Muller et al., 2010). Other DFM-specific markers are genes whose roles were described in other types of muscles, including *midline* (*mid*) (Kumar et al., 2015), *Anaplastic lymphoma kinase* (*Alk*) (Englund et al., 2003; Lee et al., 2003) and *wing blister* (*wb*) (Martin et al., 1999) that are involved in embryonic myogenesis, and *sprout* (*sty*), which regulates maturation of adult founder cells in the abdomen (Dutta et al., 2005). Concordantly, the *wb* reporter is expressed at much higher levels in DFM precursors than in IFM precursors (Figure 3E). Finally, the panel of DFM markers included *arginine kinase (Argk*), *Neurotactin* (*Nrt*), *Amalgam* (*Ama*) and *Tenascin accessory* (*Ten-a*), whose function in muscle has not been investigated. We confirmed DFM-specific expression of three of these novel markers using *Ama-GAL4* enhancer trap, GFP-tagged reporter *Argk::GFP* and anti-Nrt antibody. In each case, the wing discs were counterstained with both anti-Zfh1 and anti-Ct antibodies to visualize IFM and DFM precursors (Figure 3F-G and Figure S3B). We further showed the co-expression of Ama with Nrt, and Argk with Nrt in DFM cells, thus confirming the specificity of their expression in DFM precursors (Figure S3C).

While examining the pattern of Ama staining in the wing disc we noticed that Ama expression appears to extend into the region of IFM precursors in the anterior distal part of the prescutum (Figure S3B). To unambiguously determine which muscles Ama-expressing cells contribute to, we performed cell tracing experiments using G-TRACE (Evans et al., 2009). In this technique, GFP marks the cells that expressed *Ama* in the past (GAL4 lineage), while RFP shows the *Ama* expression at the time of the analysis (Active GAL4). In the larval wing disc, RFP expression was localized primarily to the region of DFM precursors, while GFP was broadly expressed in both IFM and DFM precursors, thus indicating that *Ama* was expressed in IFM precursors earlier in development (Figure 3H). Accordingly, both GFP-and RFP-positive cells were found in a subset of IFM precursor cells surrounding the developing IFM myotubes at 16 APF, thus suggesting that *Ama* is also expressed in developing IFM (Figure 3I). However, only GFP signal was present in mature IFM of adults. Similarly, we detected *Ama* expression in the developing DFM at 28h APF, but not in the mature DFM of adults, where only GFP was detected (Figure 3J). We conclude that *Ama* is expressed in both DFM and IFM precursors, however, only a small number of IFM precursors express *Ama* prior to the formation of myotubes, and *Ama* is expressed more broadly in DFM precursors. This is consistent with the results from scRNA-seq showing *Ama* expression in DFM clusters, but only in one IFM cluster, IFM E(spl)_1 (Figure 3B).

We conclude that DFM and IFM precursors are transcriptionally distinct and, in addition to the known markers *ct* and *vg*, these cells express a panel of muscle-specific markers during larval development. However, the expression of several DFM-specific markers can be detected in a cluster of IFM precursors, IFM E(spl)_1, albeit at a lower level.

### Exploring the heterogeneity of IFM and DFM precursors

Next, we set to determine the basis of cell heterogeneity that drives clustering of DFM and IFM precursors. The expression of the top markers for each cell population was visualized using dot plot (Figure 4A, and Supplemental Table S6). We began by examining the expression of *twist* (*twi*) and *Him* that are responsible for keeping myoblasts in an undifferentiated state and whose downregulation is required for differentiation (Anant et al., 1998; Soler and Taylor, 2009). Both *twi* and *Him* were highly expressed in two IFM clusters, IFM-E(spl)_1 and IFM-E(spl)_2, indicating that these clusters comprise undifferentiated AMPs. In contrast, IFM-E(spl)_3, IFM_1 and IFM_2 contained differentiating AMPs as *twi* and *Him* were downregulated in these cells. The expression of *twi* and *Him* is controlled by Notch (Anant et al., 1998; Bernard et al., 2006; Rebeiz et al., 2002; Soler and Taylor, 2009). Concordantly, numerous E(Spl) genes that are known targets of the Notch pathway and commonly used to access Notch activity (Zacharioudaki and Bray, 2014) were expressed in IFM-E(spl)_1 and IFM-E(spl)_2 (Figure 4A, Figure S4A). Interestingly, although E(spl) genes were still expressed in IFM-E(spl)_3, the levels of expression of both *twi* and *Him* were low, thus indicating that this cluster may represent myoblasts undergoing transition from undifferentiated to differentiated state of IFM_1 and IFM_2. Similarly, we conclude that DFM-E(spl) represents undifferentiated DFM precursors because of the high levels of expression of *twi*, *Him* and *E(spl)* genes. And, the differentiating myoblasts are grouped into a single DFM cluster, which has low *twi*, *Him* and *E(spl)* gene expression. Intriguingly, we found specific expression of *E(spl)mdelta-HLH* in DFM-E(spl), but not in IFM-E(spl) (Figure 4A), suggesting that Notch-dependent downstream response in DFM precursors may differ from IFM precursors. Additionally, we detected the expression of both *Actin 57B* (*Act57B*) and *Actin 87B* (*Act87E*) in the DFM-E(spl) cluster, which is in agreement with previous findings (Butler et al., 2003). Finally, we examined the expression of known markers of muscle differentiation, including genes encoding proteins required for myoblast fusion (*sns*, *robo*, *sli*, *blow*, *mbc*, *Vrp1*, *WASp*, *Arp2*, *rst*, *hbs*, *sing*, *Ced-12*, *Hem*), myotube attachment (*kon*, *drl*, *if*, *mys*, *Ilk*, *rhea*), sarcomerogenesis (*Act88F*, *Act79B*)), and myogenic regulators (*Mef2*, *ewg*), in IFM_1 and IFM_2. Yet, the RNA levels of these genes were too low to further delineate the differentiated state of these clusters (Supplemental Table S1).

**Figure 4.**
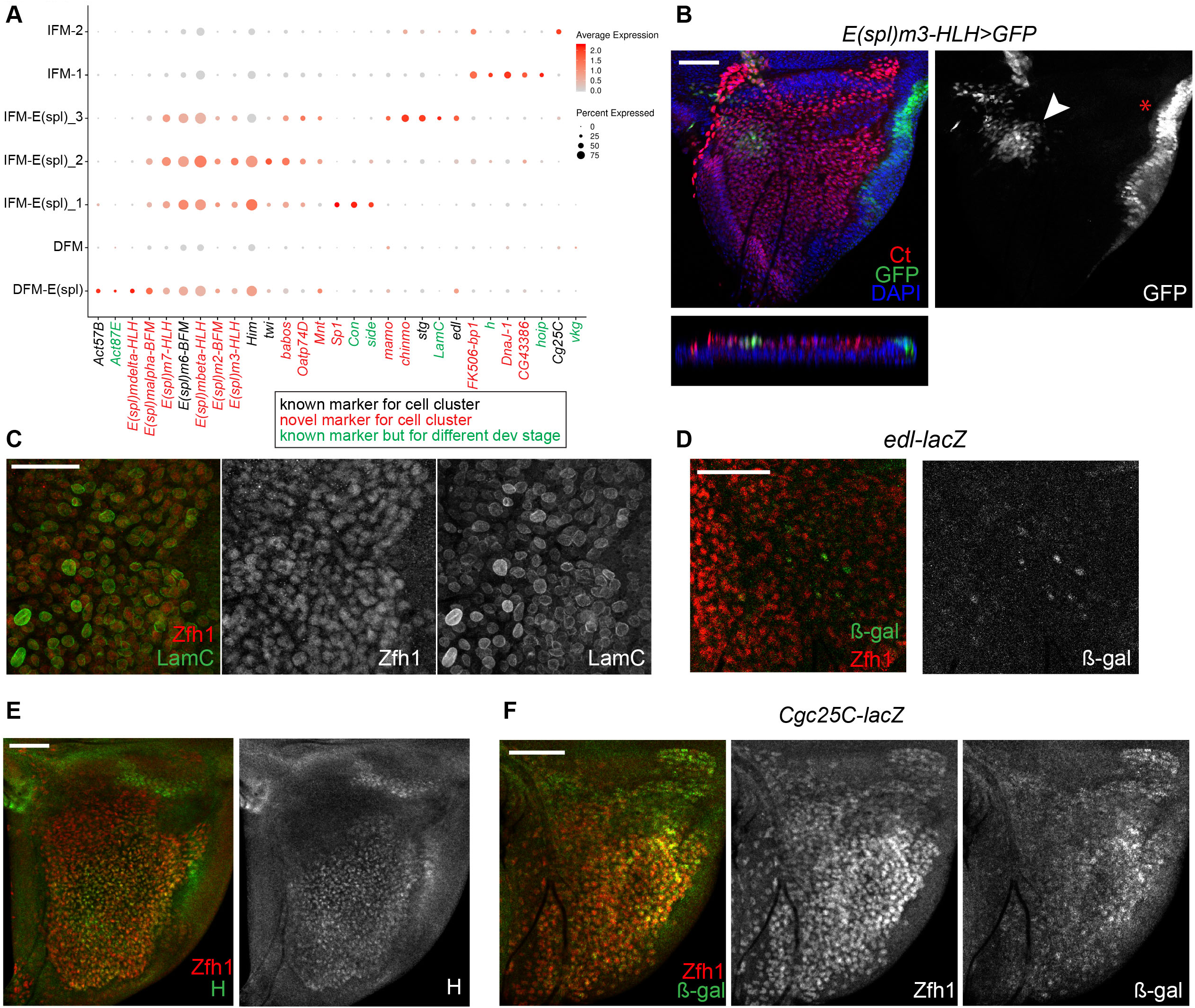
Exploring IFM and DFM precursors reveals their heterogeneity. (A) Dot plots showing the expression levels of the top variable genes between AMP clusters across 7 AMP clusters of the reference cell atlas dataset. Color intensity represents the average normalized expression level. Dot diameter represents the fraction of cells expressing each gene in each cluster Confocal single plane images of third instar larval wing discs (B) *E(spl)m3-HLH>GFP* (green) stained with anti-Ct (red). Arrowhead points to *E(spl)m3-HLH>GFP* in AMP cells. Red asterisk indicates expression of *E(spl)m3-HLH>GFP* in epithelial cells, (C) stained with anti-Laminin C (LamC, green) and anti-Zfh1 (red), (D) *edl-lacZ* stained with anti-β-gal (green) and anti-Zfh1 (red). (E) Discs stained with anti-H (green) and anti-Zfh1 (red), (F) *Cg25C-lacZ* stained with anti-β-gal (green) and anti-Zfh1 (red). Scale bars represent 50 μm. Full genotypes are (B) w-; UAS-GAL4; E(spl)m3-HLH[GMR10E12]-GAL4, (C, E) *y-, v-; +; P{CaryP}attP2*, (D) *y-, w-; edl[k06602]; +*, (F) *y-, w-; Col4a1[k00405]-lacZ; +*

The spatial expression of E(spl) genes in third larval wing disc has been extensively studied (de Celis et al., 1996b; Lai et al., 2000; Singson et al., 1994; Wurmbach et al., 1999). Interestingly, *E(spl)m6-HLH*, one of the top markers of IFM-E(spl) clusters (Figure 4A), was shown to be expressed in the specific region of the adepithelial layer near the anterior region of the presumptive lateral heminotum (Lai et al., 2000; Wurmbach et al., 1999). We used a reporter for *E(spl)m3-HLH* and found that it was expressed in a highly localized manner in the adepithelial layer (Figure 4B), and largely matching the reported pattern of *E(spl)m6-HLH* (Lai et al., 2000). Since these two genes are among the top markers of IFM-E(spl)_2 we conclude that the cells of this cluster are spatially localized to the anterior region of the presumptive lateral heminotum. The cluster IFM-E(spl)_1 was distinct from IFM-E(spl)_2 by the expression of *Sp1*, *Connectin* (*Con*) and *sidestep* (*side*) (Figure 4A), along with *Ama*, *Argk* and *wb* (Figure 3B). Both *Con* and *side* are also expressed in embryonic muscles (Nose et al., 1992). As we have described above, *Ama* was expressed in a localized manner in IFM precursors (Figure 3H and Figure S3B), which largely matches the expression pattern of *E(spl)m3-HLH*, a marker for IFM-E(spl)_1. Thus, IFM-E(spl)_1 and IFM-E(spl)_2 appear to be localized to the same region.

As described above, the cluster IFM-E(spl)_3 represented cells transitioning to the state of differentiation given the low level of *twi* and *Him*. Among other top markers for this cluster were *Chronologically inappropriate morphogenesis* (*chinmo*), *maternal gene required for meiosis* (*mamo*), *ETS-domain lacking* (*edl*), and *Lamin C* (*LamC*) (Figure 4A, Figure S4C). *LamC* is also implicated in larval muscle function and leg muscle development (Dialynas et al., 2010). We used anti-LamC antibody to examine its expression in the AMP by co-staining with anti-Zfh1. Interestingly, although LamC was expressed throughout the adepithelial layer we noticed that its expression was highly variable as some individual cells had a much higher LamC staining than others (Figure 4C). *edl-LacZ* reporter appeared to be expressed in a similar manner albeit the variation in the expression among the AMPs were not as pronounced as for LamC (Figure 4D). We conclude that IFM-E(spl)_3 comprises cells with high level of *LamC* and *edl*, and that these cells are distributed throughout the adepithelial layer unlike a highly localized position for IFM-E(spl)_1 and IFM-E(spl)_2.

Next, we examined the expression of the top markers for the two clusters of differentiating AMPs, IFM_1 and IFM_2. One of the IFM_1 markers was *hoi-polloi* (*hoip*), which is a regulator of muscle morphogenesis (Johnson et al., 2013). Another marker for IFM_1 was *h*, which is also expressed in various embryonic muscles (Fasano et al., 1988; Martin et al., 2001). As shown in Figure 4E, H was expressed throughout the adepithelial layer, but was largely excluded from the anterior region of the notum where IFM-E(spl)_1 and IFM-E(spl)_2 were located. This is in agreement with the results from our scRNA-seq indicating that *h* marks a unique AMP cluster (Figure 4A).

The cells of IFM_2 cluster expressed components of the extra cellular matrix, the type IV collagens, which is encoded by *Cg25C* and *viking* (*vkg*) genes. Both *Cg25C* and *vkg* play a role in muscle attachments in the embryo (Borchiellini et al., 1996; Hollfelder et al., 2014; Junion et al., 2007). In the wing disc, the highest expression of *Cg25C* was in the adepithelial layer located at the posterior scutellum, which was determined by co-staining wing discs with anti-Cg25C and anti-Zfh1 antibodies (Figure 4F).

In summary, scRNA-seq robustly clusters both DFM and IFM precursors based on their differentiation status. DFM precursors are grouped into DFM-E(spl) representing undifferentiated AMPs and DFM containing AMP undergoing differentiation. Notably, for IFM precursors we discerned two clusters of undifferentiated AMPs, IFM-E(spl)_1 and IFM-E(spl)_2, a cluster of cells committing to differentiation, IFM-E(spl)_3, and two clusters of differentiating AMPs, IFM_1 and IFM_2, at which the latter appears to be involved in the formation of muscle attachment sites.

### RNAi screen validates the functional importance of the novel markers identified by scRNA-seq

Among the list of specific markers for each cluster (Figure 3B, Figure 4A, Figure S5A, Supplemental Table S2, Table S3, Table S5, and Table S6) we noticed a number of novel genes that were not previously linked to the development of the adult flight muscles. To determine the role of these genes in skeletal muscle, we systematically disrupted their function exclusively in the muscle by RNAi. First, we scored viability by crossing the publicly available UAS-RNAi transgenes to the pan-muscle *Mef2-GAL4* driver (Figure 5A, Supplemental Figure S5B, Supplemental Table S7)). If no lethality was observed, adults were tested for their ability to fly as a readout of skeletal muscle function (Figure 5B, Supplemental Figure S5C). If lethality was detected at embryonic or early larval stages, then another driver, *1151*-GAL4 was used that is specifically expressed in AMP cells (Anant et al., 1998) (Figure S5D). If available, at least two RNAi line stocks targeting the same gene were tested. A total of 35 RNAi lines targeting 24 genes were screened (Supplemental Table S7).

**Figure 5.**
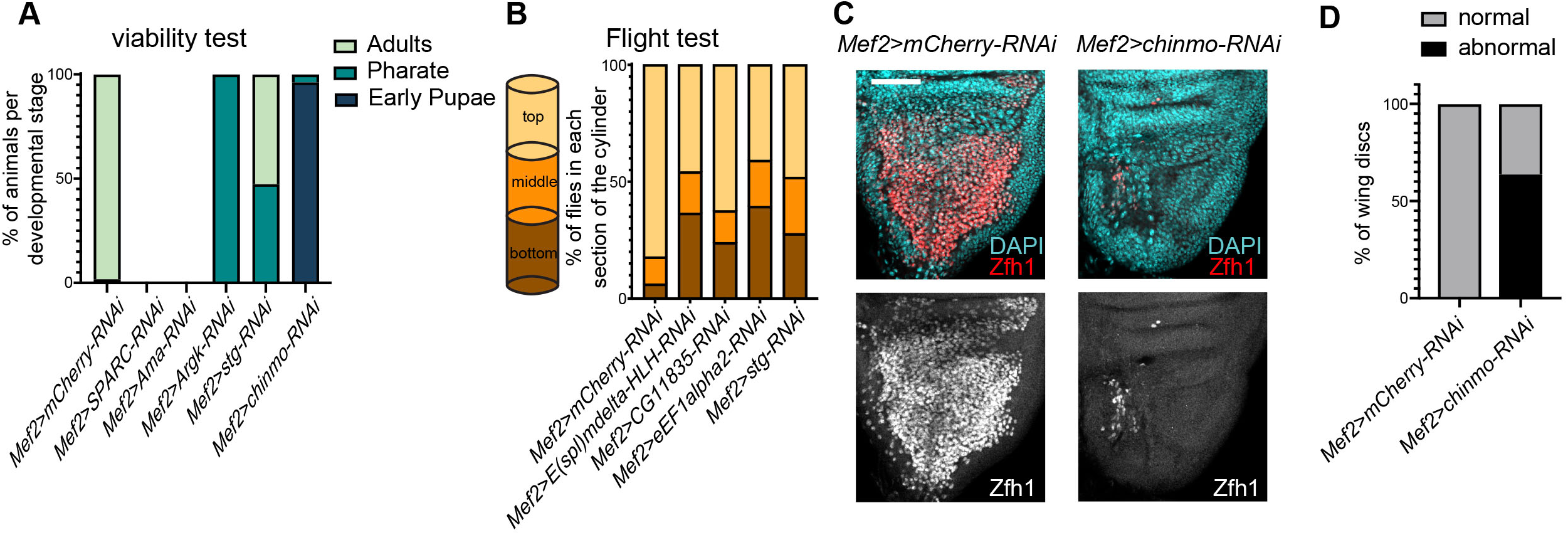
Novel markers are functionally important for the development of the flight muscles. (A) Viability test was quantified as the percentage of animals at each developmental stage relative to the number of pupae. (B) Flight ability scored as percentage of flies landing on each section of the column. (C) Confocal single plane images of third instar larval wing discs stained with anti-Zfh1 (red) and DAPI (cyan). Control (left panel) was compared to *Mef2>chinmo[HM04048]-RNAi* (right panel) to illustrate severe myoblast depletion in the knockdown. Scale bars represent 50 μm. (D) Percentage of wing discs scored as either normal or as abnormal, *i.e.* penetrant phenotype as displayed in (C). N = 32 discs, 3 independent experiments. Full genotypes are (A) *Mef2-GAL4/UAS-mCherry-RNAi, UAS-SPARC[HMS02133]-RNAi; Mef2-GAL4, Mef2-GAL4/UAS-Ama[HMS00297]-RNAi, Mef2-GAL4/UAS-Argk[JF02699]-RNAi*, *Mef2-GAL4/UAS-stg[HMS00146]-RNAi, Mef2-GAL4/UAS-chinmo[HM04048]*-RNAi (B) *Mef2-GAL4/UAS-mCherry-RNAi*, *Mef2-GAL4/ UAS-E(spl)mdelta-HLH[JF02101]RNAi*, *Mef2-GAL4/UAS-CG11835[HMS02291]-RNAi, UAS-eEF1alpha2[HMC05694]-RNAi; Mef2-GAL4, Mef2-GAL4/UAS-stg[HMS00146]-RNAi* and (C, D) *Mef2-GAL4/UAS-mCherry-RNAi* and *Mef2-GAL4/UAS-chinmo[HM04048]-RNAi*.

Seven genes scored in at least one test. One of the top candidates from the screen was *Ama*, which was expressed in both DFM and DFM-E(spl) clusters and to a lesser extent in IFM-E(spl)_1 (Figure 3B). Its inactivation using either *Mef2-Gal4* or *1151-Gal4* caused early larval and pupal lethality, respectively (Figure 5A, Figure S5D). Among the other novel genes are *Argk*, one of the top markers for DFM precursors, and *chinmo*, which is highly expressed in cluster IFM-E(spl)_3 (Figure 4A). The inactivation of these genes using *Mef2-GAL4* resulted in pupal lethality (Figure 5A). As shown in Figure 5C-D, depletion of *chinmo* by RNAi induced a severe effect on the number of AMPs in the adepithelial layer of the wing imaginal disc. However, the phenotype of *Mef2>chinmo*-*RNAi* was not fully penetrant and precluded us from further investigating the role of *chinmo*.

As expected, *E(spl)mdelta-HLH*, a DFM-E(spl) marker, scored positively (Figure 5B), which is consistent with the known role of Notch signaling in muscle development. Another hit from the screen was *string* (*stg*), whose inactivation with *Mef2-GAL4* resulted in pupal lethality (Figure 5A), while the few Mef2>*stg*-RNAi animals that survived to adulthood were flightless (Figure 5B). This is likely due to defects in the proliferation of AMP cells given a well-established role of *stg* in cell cycle. The early lethality of *Mef2>BM-40-SPARC-RNAi* (Figure 5A) is likely due to its known function in embryonic mesoderm (Martinek et al., 2002) as its inactivation with the specific adult skeletal driver *1151-GAL4* did not show muscle-related phenotype (Figure S5D). We reasoned that the inactivation of *eEF1α2* encoding the eukaryotic translation elongation factor 1 alpha 2 may exert a broad effect on protein translation, and therefore may explain why *Mef2>eEF1alpha100E* -*RNAi* animals were flightless (Figure 5B). Finally, the phenotype of *Mef2>CG11835-RNAi* was quite mild (Figure 5B).

### Amalgam is required for the expansion of the AMP pool in larval wing discs and muscle formation

We selected *Ama* for further analysis as its knockdown displayed severe muscle-related phenotype. *Ama* encodes an immunoglobulin protein that acts as a cell-adhesion molecule during axon guidance in *Drosophila* (Liebl et al., 2003). We began by profiling the proximal *1151>Ama-RNAi* wing discs by scRNA-seq. After computational processing, cell clusters were visualized using UMAP (Figure 6A). Cell type, including AMP, epithelial and tracheal cells, was assigned based on the expression of *zfh1*, *ct*, *Fas3*, *grh* and *trh* (Figure 6B) as in the reference cell atlas (Figure 1E). Overall, we have captured 3,338 cells across three replicates (Figure S6A). Strikingly, only 180 cells were AMPs among them (Figure 6A). In contrast, the wild type atlas contained 4,544 AMPs among 11,527 cells (Figure 1C), thus indicating that *Ama*-depleted AMPs are severely underrepresented. To confirm this observation, *1151>Ama-RNAi* late larval wing discs were dissected and stained with anti-Ct and anti-Zfh1 antibodies to visualize AMPs. In agreement with scRNA-seq data, there was a drastic reduction in the number of *Ama*-depleted AMP cells relative to the control (Figure 6C).

**Figure 6.**
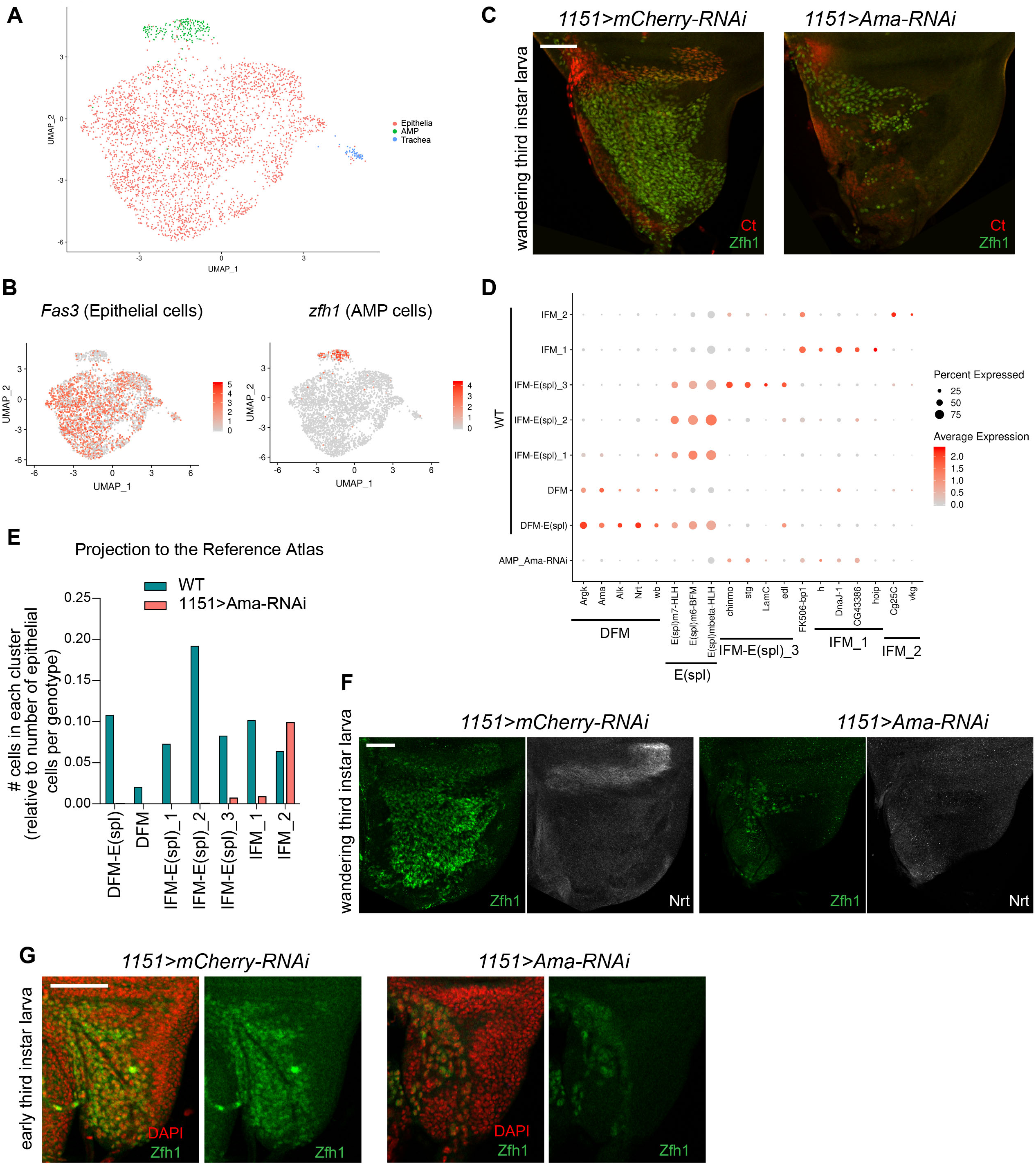
Amalgam is required for the expansion of AMP pool in larval wing disc. (A) Two-dimensional UMAP representation of single-cellRNA-seq *1151>Ama-RNAi* data set colored by cell type, including 51 tracheal, 3,107 epithelial and 180 AMP cells. (B) Average expression level of the genes *Fas3* (left panel) and *zfh1* (right panel) used as known markers to assign epithelial and AMP cells in the *1151>Ama-RNAi* dataset, respectively. (C) Confocal single plane images of *1151>mCherry-RNAi* and *1151>Ama-RNAi* wing discs at wandering third instar larval wing discs stained with anti-Ct (red) and anti-Zfh1 (green). (D) Dot plots showing the expression levels of the top markers for AMP cluster identified in reference cell atlas across the cluster of AMP cells in *1151>Ama-RNAi* and the 7 AMP clusters in control. Color intensity represents the average normalized expression level. Dot diameter represents the fraction of cells expressing each gene in each cluster (E) The AMP cells in *1151>Ama-RNAi* dataset projected into the reference single cell atlas by transferring cell type labels using Seurat. Total number of cells per cluster was normalized to the total number of epithelial cells per genotype. (F) Confocal single plane images of third instar larval *1151>mCherry-RNAi* and *1151>Ama-RNAi* wing discs stained with anti-Zfh1 (green) and anti-Nrt (white). (G) Confocal single plane images of *1151>mCherry-RNAi* and *1151>Ama-RNAi* wing discs at early third instar larval wing discs stained with anti-Zfh1 (green) and DAPI (red). Scale bars represent 50 μm. Full genotypes are *1151-GAL4;+; UAS-mCherry-RNAi* and *1151-GAL4; +; UAS-Ama[HMS00297]-RNAi*.

To further characterize *Ama*-depleted AMPs, we selected the markers for AMP clusters from the reference cell atlas (Figure 3B, 4A, Supplemental Table S8) and examined their expression in the AMP cells of *1151>Ama-RNAi* by scRNA-seq (Figure 6E). Strikingly, the expression of the DFM markers *Argk*, *Ama*, *Alk*, *Nrt* and *wb* as well as IFM-E(spl) markers, including *E(spl)m6-BFM*, *E(spl)m7-HLH and E(spl)mbeta-HLH*, was lost in *Ama*-depleted AMPs. Accordingly, the projection of *1151>Ama-RNAi* cells to the wild type reference atlas using the cell type label transfer tool in Seurat revealed that *Ama*-depleted cells were assigned to neither DFM nor IFM-E(spl) clusters (Figure 6F). In contrast, there was no significant change in the assignment of the epithelial cells of *1151>Ama-RNAi* to the reference atlas (Figure S6B) or in the expression of the epithelial marker *grh* in these cells (Figure S6C). Consistent with the loss of expression of the DFM markers in *Ama*-depleted AMPs, staining of the *1151>Ama-RNAi* wing discs with antibodies against the DFM marker Nrt and the marker Zfh1 revealed the complete lack of DFM precursors (Figure 6G).

Although the majority of the *Ama-*depleted AMP cells were assigned to IFM_2 cluster (Figure 6F) several IFM_2 markers, including *Cg25C* and *vkg*, were misexpressed (Figure 6E), implying that the transcriptional program in *Ama*-depleted AMPs is dramatically different from the wild type AMPs. Interestingly, *Ama*-depleted AMPs no longer express E(spl) genes, thus indicating the loss of the signal to maintain undifferentiated state. To determine whether *Ama*-depleted AMPs prematurely induce differentiation program, we examined the expression of known differentiation markers, including genes involved in myoblast fusion and muscle attachment (Figure S6D). However, for the exception of *blow* and *mys*, the differentiation markers were not induced indicating that the *Ama*-depleted AMPs do not undergo premature differentiation.

The reduction in the number of the *Ama*-depleted AMPs (Figure 6C) could be due to either an increased apoptosis or a reduction in cell proliferation. To decipher between these two possibilities, the wing discs were stained with anti-Dcp1, to monitor apoptotic cells, and anti-PhosphorHistone H3 (pH3), a marker of mitotic cells. The depletion of *Ama* appears to affect cell proliferation as no pH3 positive cells were observed among *Ama*-depleted AMPs, while the level of apoptosis was undetectable (Figure S6E-F). We acknowledge that the low number of AMPs in *1151>Ama-RNAi* precludes accurate quantification of the phenotypes. Notably, the number *Ama*-depleted AMPs remained reduced in early third instar larval wing disc indicating that upon depletion of *Ama* the AMP cells fail to undergo a massive expansion that normally occurs at this stage (Figure 6G) (Gunage et al., 2014).

To determine whether the *Ama*-depleted AMPs are competent to form muscles, we examined the formation of DFM and IFM in *1151>Ama-RNAi* at early stages of pupal development. Although AMPs properly migrated and were detected at early pupae stages in *1151>Ama-RNAi*, no developing DFM were visible at 40 h APF (Figure 7A). Accordingly, no developing IFM were detected in *1151>Ama-RNAi* at 16 h APF (Figure 7B) nor at 20 h APF (Figure S7). Notably, larval oblique muscles, which are used as templates for IFM formation, were not found, suggesting that these may have not escaped histolysis.

**Figure 7.**
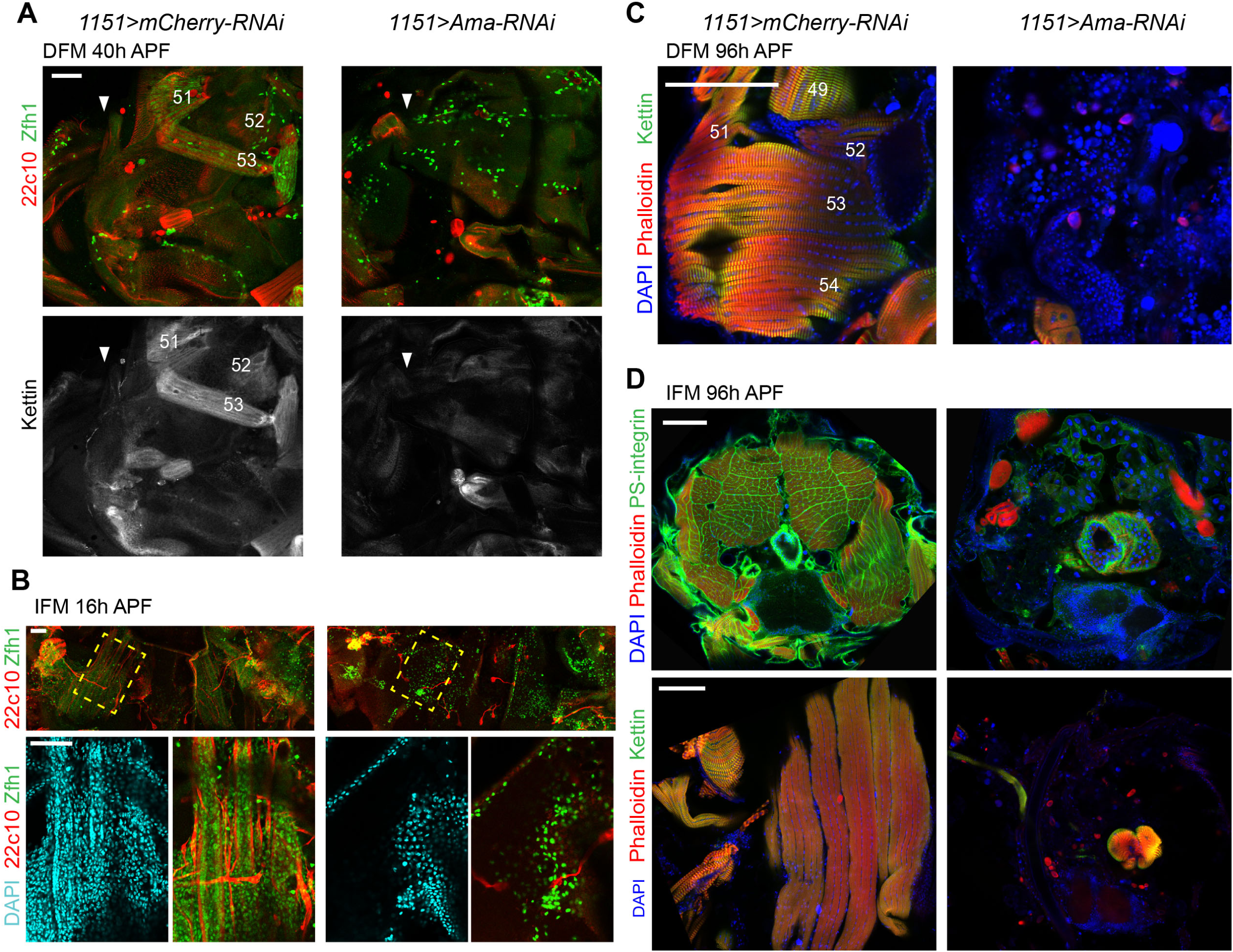
The loss of Amalgam impairs the formation of both IFM and DFM. Confocal single plane images of DFM (A, C) and IFM (B, D) at 40h APF (A), 16 h APF (B), and pharate (C, D) of *1151>Ama-RNAi* animals compared to control *1151>mCherry-RNAi*. (A) Forming DFM at 40 h APF stained with anti-Zfh1 (green), anti-Futsch/22c10 (red) and anti-Kettin (white). White arrows point to the wing hinge, wings pointing left, anterior up. (B) Forming IFM (DLM) at 16 h APF, stained with anti-Futsch/22c10 (red), anti-Zfh1 (green) and DAPI (cyan). Yellow-dashed box indicates magnified area (bottom panel). Anterior up. (C) DFM at 96 h APF stained with Phalloidin (red), anti-Kettin (green) and DAPI (blue). Anterior up, dorsal right. (D) 96 h APF thorax, transverse section (top) stained with anti-PS-integrin (green), Phalloidin (red) and DAPI (blue), dorsal up; sagittal section (bottom) stained with anti-Kettin (green), Phalloidin (red) and DAPI (blue), anterior up, dorsal right. DFM are numbered in white as in (Lawrence, 1982) Scale bars are 50 μm (A, B), and 100 μm (C, D). Full genotypes *are 1151-GAL4; +; UAS-mCherry-RNAi* and *1151-GAL4; +; UAS-Ama[HMS00297]-RNAi*.

To exclude the possibility that *Ama*-depleted AMPs are simply delayed in development and form muscle later, we examined the adult skeletal muscles in *1151*>*Ama RNAi* pharate at 96 h APF. Thoracic sections were stained with Phalloidin and antibodies against Kettin and PS-Integrin to visualize the myofibril structure of DFM and IFM by immunofluorescence. Strikingly, DFM were completely missing in *1151>Ama-RNA*i in comparison to control animals (Figure 7C). Likewise, IFM muscles were detected in neither transverse nor sagittal thoracic sections (Figure 7D, top and bottom panels, respectively).

From these data, we conclude that *Ama*-depleted AMPs are severely impaired in their proliferation capacity. Additionally, the transcriptional program of the remaining *Ama*-depleted AMPs is significantly perturbed as they completely fail to form flight muscles.

## Discussion

The AMP cells associated with the larval wing discs give rise to two distinct types of adult flight muscles, the fibrillar IFM and the tubular DFM, with distinct physiology, size, contractile properties, and metabolic characteristics. To account for these differences, the AMPs undergo complex diversification during development that culminates in the expression of fiber type-specific structural genes, among others. However, it is not clear when the major differences in their intrinsic transcriptional programs arise. With the advancement in single-cell technologies, it is now feasible to dissect the transcriptomes of individual cells and accurately identify cell types, cell states, gene signature, and major genetic drivers of developmental programs. In this study, we performed single cell RNA-seq to build a reference atlas of 4,544 AMPs associated with the third instar larval wing disc. The atlas represents approximately 1.8x cellular coverage based on the estimated number of 2,500 AMPs per wing disc at this developmental stage (Gunage et al., 2014). By querying the cell atlas, we dissected cell heterogeneity between AMPs and explored early events in establishing muscle diversity. Here, we report three main findings.

First, we show that in the third instar larva the IFM and DFM precursors have distinct transcriptional programs. Our data are in agreement with previous studies showing that the divergence of the AMP cells correlates with the differential expression of *vg* and *ct* (Sudarsan et al., 2001). It has been suggested that Wg emanating from the wing disc epithelium is required for the maintenance of Vg expression, and thus helps to spatially subdivide AMPs into DFM and IFM precursors (Sudarsan et al., 2001). One implication of our results is that the divergence is not merely because DFM and IFM precursors are in distinct regions of the notum defined by the pattern of Wg expression, but is rather due to the extensive differences in the transcriptional programs between IFM and DFM precursors that are not limited to *vg* and *ct*. Interestingly, such transcriptional changes occur prior to the expression of fiber-specific genes that are thought to distinguish fiber types at the molecular level (Schiaffino and Reggiani, 2011). At least two distinct regulatory mechanisms mediated by Salm, Exd and Hth, govern fiber-specification fate by regulating the expression of components of myofibrillar structure in late myogenesis (Bryantsev et al., 2012; Schönbauer et al., 2011). The differences in gene expression in the proliferating AMPs that we report here raise the question of what other factors are contributing to the divergence of muscle diversity at early stages of myogenesis.

Our work identifies a number of cell type-specific markers that distinguish DFM precursors from IFM precursors, and were largely missed in previous studies using bulk RNA-seq (Spletter et al., 2018; Zappia et al., 2019). We acknowledge that this list is likely incomplete due to a low depth of sequencing in Drop-seq. Interestingly, the list of markers includes *trol*, *nkd*, *Alk*, *sty*, *wb*, *mid*, *Con*, *side*, *LamC*, *h*, *hoip*, and *vkg*, which are also involved in the formation of embryonic muscles, thus indicating that these genes are reused again during the development of adult skeletal muscle.

Since the AMPs are associated with the epithelial cells of the wing disc, we also recovered about 7,000 cells of epithelial cells and 270 tracheal cells. Although this was not the goal of this work, we mapped 17 epithelial cell clusters to the wing disc fate map. Together with two recent studies (Bageritz et al., 2019; Deng et al., 2019), our work provides a valuable resource for the Drosophila community as a part of The Fly Cell Atlas initiative (https://flycellatlas.org/).

The second conclusion of our work is that the populations of IFM and DFM precursors are highly heterogeneous as we identified five IFM and two DFM clusters that represent cells at various states of differentiation. Consistent with the well-documented role of Notch pathway in the formation of IFM, we found clusters of AMPs expressing *E(spl)* genes, which are indicative of Notch activity, and clusters with low expression of *E(spl)* which are, likely, more differentiated AMPs. Interestingly, the expression of some downstream effectors of Notch pathway, such as *E(slp)-mdelta*, differs between the IFM and DFM precursors, thus suggesting that although Notch is the main driving force in the regulatory network, the output is likely modulated by other genetic cues that depend on each type of AMP. Noteworthy, the expression of *E(spl)m3* (this work) and *E(spl)m6* (Lai et al., 2000), which are the top markers for the clusters IFM-E(spl)_1 and IFM-E(spl)_2, is localized within the adepithelial layer near the anterior region of the presumptive lateral heminotum. These data suggest that, at least, the populations IFM-E(spl)_1 and IFM-E(spl)_2 are spatially restricted. This is in contrast to other clusters, such as IFM-E(spl)_3, which are largely distributed throughout the adepithelial layer. Curiously, in addition to *E(spl)* genes, the cluster IFM-E(spl)_1 expresses several DFM markers, such as *Ama*, *Argk*, and *wb*. We disfavor the explanation that the cluster IFM-E(spl)_1 contains cell doublets of IFM and DFM precursors as this cluster shows high *vg* and low *ct* expression, an IFM hallmark (Sudarsan et al., 2001). Furthermore, we confirmed the expression of *Ama* in IFM precursors by genetic tracing experiments.

Third, our work provides a framework for leveraging scRNA-seq to identify novel genes that are functionally important in a particular biological process. Concordantly, several novel muscle genes identified here as markers for AMPs also scored in a large-scale screen for genes involved in muscle morphogenesis and function (Schnorrer et al., 2010). Using the list of marker genes as an entry point in an RNAi screen we discovered *Ama*, whose inactivation caused severe muscle defects. Ama was shown to act as a ligand for the cell-adhesion molecule Nrt during axon guidance in *Drosophila* embryogenesis (Fremion et al., 2000). It has been proposed that Ama facilitates Nrt-mediated adhesion by functioning as a linker between two Nrt-expressing cells. Intriguingly, both Nrt and Ama are highly expressed in the DFM precursors raising the possibility that Ama may similarly interact with Nrt in the AMPs to regulate adhesion. We found that the depletion of Ama led to a severe reduction in the number of AMPs, which is most likely due to the failure to undergo expansion during larval stages. Interestingly, Ama was shown to regulate the expression of Cyclin E through Hippo pathway (Becker et al., 2016), which may explain the proliferative defects we report here. Strikingly, the remaining *Ama*-depleted AMPs are unable to form myotubes, which is likely caused by the highly abnormal transcriptional profile of the *Ama*-depleted AMPs, as revealed by scRNA-seq. Thus, the failure to properly execute the myogenic transcriptional program may underlie muscle defects upon the loss of Ama function.

One limitation of this approach is that it may fail to identify low expressed genes or conversely the UAS-RNAi lines may not be robust enough to display a phenotype with the specific-muscle driver. Therefore, the list of candidates that we report here is likely to be incomplete. Nevertheless, this strategy can be applied to other scRNA-seq datasets to address the functional significance of novel marker genes identified as a part of scRNA-seq computational pipeline. Collectively, our work illustrates the power of combining single cell genomics with genetic approaches to address important questions in developmental biology.

## Methods

### Fly maintenance and stocks

All lines used here are listed in Supplemental Table S9. All fly crosses were kept at 25°C in vials containing standard cornmeal-agar medium.

### Fly viability assay

Adult flies able to eclose from the pupal case, dead pharate, and dead pupae, were counted to score the percentage of flies that made it to each developmental stage. Pupal developmental stages were determined by observing biological markers of metamorphosis. At least 52 flies per genotype were screened in total (unless lethal at early stages), with a minimum of two experimental replicates.

### Flight test

Males no older than a week were collected on CO2 and kept at 25°C for at least 24 hours for recovery, and then at room temperature for another hour for acclimation. Flies were then flipped into a 2L graduated cylinder lined with a 432 mm high piece of paper coated in mineral oil. Flies landed on the paper at different heights depending on their ability to fly. A picture of the unfurled paper was then taken, and the landing spot for each fly was unbiasedly scored as one of the three sections (top, middle or bottom) by an ImageJ plugin (script available upon request). Frequencies were then analyzed and plotted. Experiments were carried out at least twice for genotypes presenting a phenotype, at least 25 flies per genotype (except for *Mef2>midHMC03082-RNAi*, n=15).

### Dissection and immunofluorescence

#### Imaginal wing discs

larvae were dissected in Phosphate Buffered Saline (PBS) pH 7.4 and immediately fixed with 4% formaldehyde in PBS for 30 minutes (15 min whenever anti-Ct antibody was used), permeabilized twice in 0.3% Triton X-100-PBS for 10 min and blocked in 10% Normal Donkey Serum (NDS) 0.1% Triton X-100-PBS for 1 hour. Samples were incubated in primary antibodies overnight at 4°C in 10% NDS 0.1% Triton X-100-PBS, washed for 5 min three times with 0.1% Triton X-100-PBS, then incubated for 1 hr with secondary antibodies and dyes in 10% NDS 0.1% Triton X-100-PBS. Samples were washed with 0.1% Triton X-100-PBS five times for 5 min.

#### Developing pupal flight muscles

thoraces were dissected as in (Weitkunat and Schnorrer, 2014), fixed for 15 min with 4% formaldehyde in PBS, permeabilized in 0.3% Triton X-100-PBS three times for 20 min and blocked in 10% NDS 0.1% Triton X-100-PBS for 120 min. Then primary antibody was incubated overnight at 4°C in 10% NDS 0.1% Triton X-100-PBS, four 25 min washes in 0.1% Triton X-100-PBS, secondary antibody was incubated for 120 min in 10% NDS 0.1% Triton X-100-PBS, and four 15 min washes in 0.1% Triton X-100-PBS.

#### Adult and pharate flight muscles

For transverse sections, flies were snap-frozen in liquid nitrogen, cut twice with a razor and fixed for 2 h in 4% formaldehyde in relaxing buffer (20mM phosphate buffer, pH 7.0; 5 mM MgCl2; 5 mM EGTA). For the sagittal IFM and DFM sections, thoraces were cut from the animals, incubated in relaxing buffer for 15 min, fixed for 30 min in 4% formaldehyde in relaxing buffer, cut through the appropriate sagittal plane with a Sharpoint 22.5° Stab Knife (ID#72-2201), and then fixed for an additional 15 min as described in (Schnorrer et al., 2010). All subsequent solutions contained 0.3% Triton X-100 for transverse and sagittal IFM and 0.5% Triton X-100 for DFM. Four 15 min washes for transverse and three 10 min for sagittal sections, 2-hour incubations in blocking solution (PBS + 2% Bovine Serum Albumin (BSA) + Triton 100-X). Sections were incubated with primary antibodies in blocking solution overnight at 4°C, washed four times with PBS solutions, 15 min for transverse and 10 min for sagittal sections. Secondary antibody incubations were 2 h long in 0.3% Triton X-100 10% NDS-PBS for transverse and sagittal IFM, and 0.5% Triton X-100 2% BSA in PBS for DFM sections. Finally, all sections were washed four times for 10 min.

Primary antibodies, secondary antibodies and dyes used in this work are listed in Supplemental Table S9. After washing off the secondary antibodies, all samples were stored in 0.5% propyl gallate 50% glycerol at 4°C until mounted on glass slides.

### Microscopy

All images were taken with a Zeiss LSM Observer.Z1 laser-scanning confocal microscope, using 10x/0.30, 20x/0.8, 40x/1.20, 63x/1.40, 100x/1.45 objectives. Pinhole was kept at 1 AU, laser and gain was kept consistent within experiments (e. g. keeping the same settings for control and knockdown). For z-stacks, the optimal slice size suggested by the microscope software was used. Orthogonal views were taken from the z-stacks using ImageJ. Only one representative image per experiment is shown.

### Software

The software used was: NIH ImageJ 1.52k5 https://imagej.nih.gov/ij/ for image visualization, Adobe Photoshop CS6 version 13.0.6 https://www.adobe.com/products/photoshop.html for image editing, GraphPad Prism version 8.0.1 https://www.graphpad.com/scientific-software/prism/ to create all graphs, Adobe Illustrator CC 23.0.2 https://www.adobe.com/products/illustrator.html for figure editing.

### Single-cell preparation

Tissue was dissociated into single cell suspension as in (Ariss et al., 2018). Briefly, wandering third instar larvae were harvested. Wing discs were dissected in cold PBS1x, pouch was manually removed with microblade. Notum and hinge were collected and processed for dissociation in a final concentration of 2.5 mg/mL Collagenase (Sigma #C9891) and 1x trypsin (Sigma #59418C) in Rinaldini solution. The microcentrifuge tube was horizontally positioned on a shaker set at 225 rpm for 20 min at RT. Cells were washed twice and resuspended in 0.04% BSA-PBS. Cell viability and concentration were assessed by staining cells with Trypan blue and counting using a hemocytometer.

### Sample preparation for scRNAseq

For Drop-seq we followed the protocol version 3.1 (12/28/15) as in (Macosko et al., 2015b) posted in http://mccarrolllab.org/dropseq/ with the following modifications. The lysis buffer contained 0.4% Sarkosyl (Sigma). The number of cycles in the PCR step post exonuclease are 4, and then 12. The cDNA Post PCR was purified twice with 0.6x Agencourt AMPure XP (Beckman Coultier). The tagmented DNA for sequencing was purified twice: first using 0.6x and the second time using 1x Agencourt AMPure XP (Beckman Coultier).

### High-throughput sequencing

Quality control of both amplified cDNA and sequencing-ready library was determined using Agilent TapeStation 4200 instrument. All Drop-seq libraries were sequenced on a NextSeq instrument (Illumina). Sequencing was done at the University of Illinois at Chicago Sequencing Core (UICSQC).

### Preprocessing Raw Datasets of scRNAseq

The Drop-seq samples were processed for read alignment and gene expression quantification following Drop-seq cook-book (version 1.2 Jan 2016)7 (http://mccarrolllab.com/dropseq/) (Macosko et al., 2015b). We used STAR aligner to align the reads against *Drosophila melanogaster* genome version BDGP6 (Ensembl version 90). Quality of reads and mapping were checked using the program FastQC (https://www.bioinformatics.babraham.ac.uk/projects/fastqc/). Digital Gene Expression (DGE) matrix data obtained from an aligned library was done using the Drop-seq program DigitalExpression (integrated in Drop-seq_tools-1.13). Number of cells that were extracted from aligned BAM file is based on knee plot which extracts the number of reads per cell, then plot the cumulative distribution of reads and select the knee of the distribution.

### scRNA-seq data analysis

The packages Seurat (version 3.0.0) and R (version 3.5.3) were used to analyze datasets and to generate plots. We followed the standard tutorial instructions from the Seurat website (https://satijalab.org/seurat/) (Stuart et al., 2019). First, the gene expression matrices were subjected to an initial quality control analysis. Low-quality cells were filtered out using 200 and 2,500 gene/cell as a low and top cutoff, respectively, and min.cell = 5. Additionally, markers of cellular stress, such as content of mitochondrial, heat shock protein and 28sRNA, were set to less than 10, 6, and 1.5% reads per cell, respectively. These four variables were regressed out when scaling. An integrative analysis was done to generate the reference cell atlas for the wild type wing discs by determining the anchor points in each dataset using Seurat pipeline. The top 2000 variables genes were used as input for PCA analysis to run linear dimensional reduction. Cell were clustered using a graph-based method. The first 30 principle components were selected to run non-linear dimensional reduction (UMAP) using granularity 2.5 for reference atlas and 1.5 for *1151>Ama-RNAi* dataset. Clusters were visualized with UMAP. Clusters of cells that were not evenly distributed among replicates, i.e. more than 50% bias for one sample, which indicates a batch effect between samples, were removed from the analysis, and then dataset was reanalyzed. Also, the number of genes, and content of mitochondrial, rRNA and heat shock protein among the clusters was examined across the clusters. The analysis was performed only on the cell barcodes as listed in Supplemental Table S10. The dimensions of the final datasets are 11,874 genes by 11,527 cells for the reference atlas, and 8,483 genes by 3,338 cells for Ama-knockdown.

Cluster-specific markers were determined by calculating differential expression using non-parameteric Wilcoxon rank sum test. In the reference atlas, two IFM-E(spl) clusters originally showed same top markers. Thus, to prevent an artifact of overclustering, these two clusters were then labeled as IFM-E(spl)_2.

### Data availability

Dropseq scRNA-seq data have been deposited in the NCBI Gene Expression Omnibus database (GEO, https://www.ncbi.nlm.nih.gov/geo/) and are accessible through the accession number GSE138626 (https://www.ncbi.nlm.nih.gov/geo/query/acc.cgi?acc=GSE138626)

## Acknowledgements

We thank T.V. Orenic for helping with the assignment of the epithelial clusters, R.M. Cripps, F. Schnorrer, M. Spletter for helpful discussions, and Anna Barque for helping with crosses and staining. We are grateful to R.M. Cripps, the Bloomington Drosophila Stock Center (supported by NIH grant P40OD018537), the Vienna Drosophila Resource Center, and the TRiP at Harvard Medical School for fly stocks, the Babraham Institute, and the Developmental Studies Hybridoma Bank (DSHB) for antibodies; and to Flybase for online resources on the Database of Drosophila Genes & Genomes. We thank the University of Illinois at Chicago Sequencing Core (UICSQC) for sequencing. This work was supported by NIH grant R35GM131707 (to M.V.F.)

## Author contributions

M.P.Z. and M.V.F. conceived the project, designed the experiments, analyzed data, and wrote the manuscript. M.P.Z. analyzed all the single-cell RNA-seq datasets. L.d.C. generated all immunofluorescences, the data from the screen and helped M.P.Z. with figure preparation. M.M.A. performed Drop-seq on the wild type wing discs. A.B.M.M.K.I. helped with the bioinformatics data analysis of high-throughput sequencing data sets. All authors reviewed and edited the manuscript.

