## Supplemental Information for "A single-cell transcriptome atlas of the adult muscle precursors uncovers early events in fiber-type divergence in *Drosophila*"

#### Supplemental Figures

##### Supplemental Figure S1. Related to Figure 1

(A) UMAP plot of reference single cell atlas generated using single-cell RNA-seq from third instar larval wing discs colored by replicates (A) and split by replicates (B). Eight wild type samples were used. Four replicates *1151-GAL4* are indicated as 1151 and 4 replicates *1151>mCherry-RNAi* are labeled as Cherry.

(C) Total number of cells per sample used in reference atlas after applying filters and removing clusters that were unevenly represented across replicates.

(D) Dimensions of the data was reduced by PCA. Genes (rows) and cells (columns) were ordered by their PCA scores. Heatmap showing distribution of the 500 most extreme cells and 30 most extreme genes. PC1 separates epithelial cells from AMP cells as indicated by AMP genes, such as *Him*, *zfh1*, and epithelial genes, including *Fas3* and *grh*.

(E) Confocal single plane images of wandering third-instar larval wing discs and orthogonal views of the discs illustrating the layering and distribution of AMPs and epithelial cells within the notum, *twi-lacZ* stained with anti-β-galactose (β-gal, red) for AMPs and GFP-tagged grainy head (*grh::GFP*, green) for epithelial cells. Scale bars represent 50 μm. Full genotype is *w-; twi-lacZ; grh[VK00033]::GFP*

(F) Dot plot showing the expression of the main markers distinguishing Trachea_1 from Trachea_2 across all clusters of the reference atlas. Color intensity represents the average normalized expression level. Dot diameter represents the fraction of cells expressing each gene in each cluster.

**Supplemental Figure S2. Related to Figure 2**

(A) Average expression level of the genes used as known markers to map localization of clusters to the cell fate map of the wing discs in the epithelial clusters assigned as in the reference atlas dataset. Marker gene for Epi_5 and 9 is *CG17278*, Epi_13 is *caup*, *ara*, *mirr* and *vn*, Epi_2 and 3 is *knrl*, Epi_15 is *tup*, and PM_2 is *Antp*.

(B) Violin plot representing the distribution of expression levels of *scabrous* (*sca*) in the epithelial clusters assigned as in the reference atlas dataset.

**Supplemental Figure S3. Related to Figure 3**

(A) Heatmap showing the 10 tops differentially expressed gene markers for each AMP cluster of the reference single cell atlas dataset. Cells (columns) are clustered by the expression of the main marker genes (rows).

(B, C) Confocal single plane images of third instar larval wing discs (B) *Ama>GFP* (green) stained with anti-Ct (red) and anti-Zfh1 (cyan), (C) magnification of DFM precursor area in *Ama>ChFP* and *Argk::GFP* stained with anti-Nrt (red).

Scale bars represent 50 μm. Full genotypes are (B) *w-; UAS-GFP; Ama[NP1297]-GAL4*, (C) *w-; +; Ama[NP1297]-GAL4/UAS-mCD8.ChRFP* and *y-, w-; +; Argk[CB03789]::GFP.*

**Supplemental Figure S4. Related to Figure 4**

(A-E) Average expression level of the genes identified as markers for the AMP clusters assigned as in the reference atlas dataset. (A) Marker gene for IFM-E(spl) and DFM-E(spl) are *E(spl)-m6-BFM*, *E(spl)-m7-HLH, E(spl)-m3-HLH* (B) IFM-E(spl)_1 is *Sp1*, (C) IFM-E(spl)_3 are *LamC* and *edl*, (D) IFM_1 is *h* and (E) IFM_2 is *Cg25C*.

**Supplemental Figure S5. Related to Figure 5**

(A) Dot plot showing the expression of the differentially expressed genes across AMP clusters of the reference single cell atlas. Color intensity represents the average normalized expression level. Dot diameter represents the fraction of cells expressing each gene in each cluster

The muscle-specific (B, C) *Mef2-GAL4* and (D) *1151-GAL4* drivers were crossed to *UAS-RNAi* lines to knockdown the expression of the differentially expressed gene markers for the AMP clusters. (B, D) Viability test was assessed by scoring the percentage of viable flies at early pupa, pharate and adult stages. (C) Flight ability was scored as the percentage of flies landing on each section of the column (top, middle and bottom).

**Supplemental Figure S6. Related to Figure 6**

(A) UMAP plot of *1151>Ama-RNAi* using single-cell RNA-seq from third instar larval wing discs colored by cell type, including AMP, epithelial and tracheal cells, and split by replicate. Three samples were used. Total number of cells after applying filters were 1232, 706 and 1400 for Rep1, Rep4 and Rep5 respectively.

(B) The cells from *1151>Ama-RNAi* dataset were projected into the reference single cell atlas. Number of cells per cluster relative to the total number of epithelial cells in each genotype is plotted across epithelial cells.

(C) Violin plot representing the distribution of expression levels of the epithelial marker *grh* in all the clusters split by *1151>Ama-RNAi* and reference cell atlas dataset.

(D) Dot plots showing the expression levels of the known differentiation genes across the cluster of AMP cells in *1151>Ama-RNAi* and the reference cell atlas. Color intensity represents the average normalized expression level. Dot diameter represents the fraction of cells expressing each gene in each cluster

(E-F) Confocal single plane images of third instar larval *1151>mCherry-RNAi* and *1151>Ama-RNAi* wing discs stained with anti-Ct (green), DAPI (red) and (E) anti-pH3 (magenta) or (F) anti-Dcp1 (magenta).

Scale bars represent 50 μm. Full genotypes are *1151-GAL4;+; UAS-mCherry-RNAi* and *1151-GAL4; +; UAS-Ama[HMS00297]-RNAi.*

**Supplemental Figure S7. Related to Figure 7**

Confocal single plane images of *1151>mCherry-RNAi* and *1151>Ama-RNAi* forming IFM (DLM) at 20 h APF stained with anti-Zfh1 (green), anti-Futsch/22c10 (red) and DAPI (cyan). Scale bar represents 50 μm. Anterior up. Full genotypes are *1151-GAL4; +; UAS-mCherry-RNAi* and *1151-GAL4; +; UAS-Ama[HMS00297]-RNAi.*

**Supplemental Tables**

**Supplemental Table S1. Average expression of all genes across all cluster found in the reference single-cell atlas. Related to Figure 1**

List of average gene expression across all clusters.

**Supplemental Table S2. Markers for each cluster from the reference single-cell atlas. Related to Figure 1**

List of all genes differentially expressed across all clusters. Only positive markers with threshold set p_val_adj<0.05 for all comparisons.

**Supplemental Table S3. Markers for epithelial, tracheal and AMP cells from the reference single-cell atlas. Related to Figure 1**

Cell type was assigned to each cluster based on the expression of *zfh1*, *trh* and *Fas3*, then clusters were grouped in three cell types and markers for each cell type were found. List of all genes differentially expressed between cell type, threshold set p_val_adj<0.05 for all comparisons. Data is shown in each tab as Epithelia markers, Trachea markers, AMP markers.

**Supplemental Table S4. Markers for Trachea_1 compared to Trachea_2 were analyzed in the reference single-cell atlas. Related to Figure S1**

List of all genes differentially expressed between clusters, threshold set p_val_adj<0.05, sorted by avg_logFC.

**Supplemental Table S5. Markers for DFM precursors and IFM precursors in the reference single-cell atlas. Related to Figure 3.**

List of all genes differentially expressed between IFM and DFM precursors, threshold set p_val_adj<0.05 sorted by avg_logFC. Cell type was assigned to each cluster based on the differential expression of ct and vg, then clusters were grouped in two AMP precursor types and markers for each AMP type were found.

**Supplemental Table S6. Markers for each AMP cluster in the reference single-cell atlas. Related to Figure 4.**

List of all genes differentially expressed between AMP clusters, threshold set p_val_adj<0.05 for all comparisons, sorted by avg_logFC. Pairwise comparisons are as follows:

(a) DFM-E(spl) vs DFM,

(b) IFM-E(spl) vs IFM. The three clusters IFM-E(spl)_1, IFM-E(spl)_2 and IFM-E(spl)_3 were grouped and compared to the group IFM_1 and IFM_2,

(c) IFM_1 vs IFM_2,

(d) IFM-E(spl)_1 vs grouped IFM-E(spl)_2 and IFM-E(spl)_3

(e) IFM-E(spl)_2 vs grouped IFM-E(spl)_1 and IFM-E(spl)_3,

(f) IFM-E(spl)_3 vs grouped IFM-E(spl)_1 and IFM-E(spl)_2.

**Supplemental Table S7. Functional analysis of AMP markers by screening muscle-related phenotype. Related to Figure 4.**

The *UAS-RNAi* lines were crossed to (a) *Mef2-GAL4* and to (b) *1151-GAL4* drivers if cross to *Mef2-GAL4* showed early lethality. List of stocks and genes used in the screen. Both viability and flight ability were assessed unless phenotype was lethal.

**Supplemental Table S8. Average expression of all genes across AMP cluster of combined datasets of *1151>Ama-RNAi* and WT (reference atlas). Related to Figure 6**

List of average gene expression across AMP clusters.

**Supplemental Table S9. List of reagents. Related to Methods**

List of reagents, including (A) fly stocks and (B) antibodies, used in this work.

**Supplemental Table S10. List of cell barcodes. Related to Methods**

List of cell barcodes used in the (A) the reference atlas and (B) *1151>Ama-RNAi* dataset after filters were applied.

Sup Fig_S1

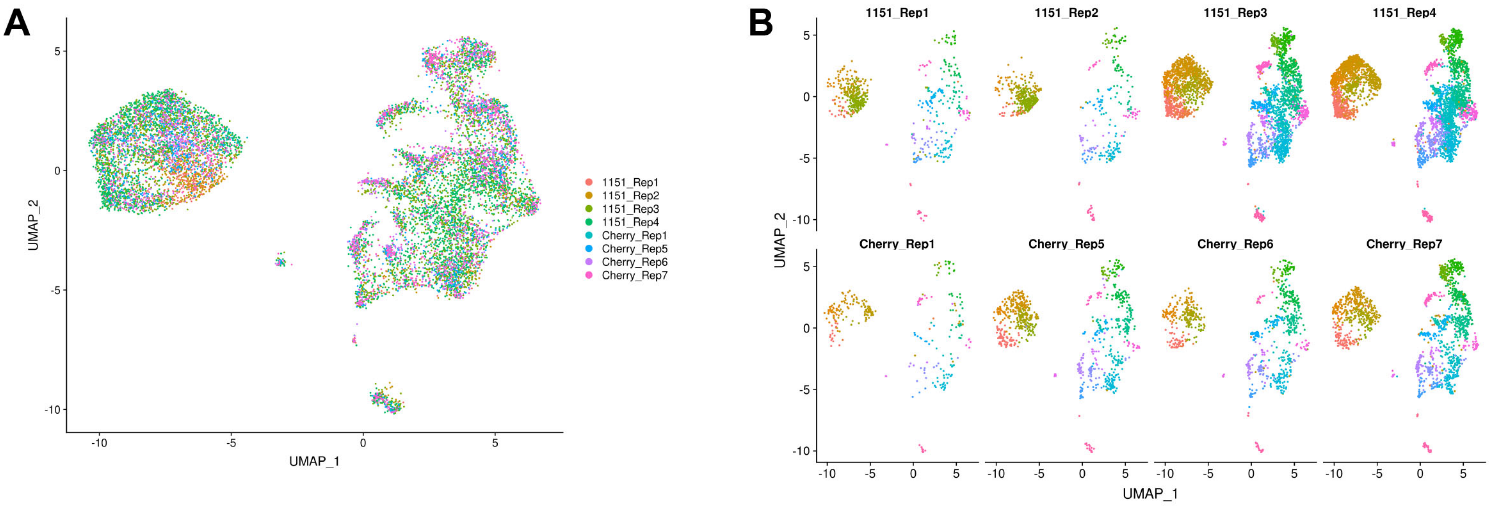


C

| Replicate | Number of cells |
|---|---|
| 1151_Rep1 | 611 |
| 1151_Rep2 | 632 |
| 1151_Rep3 | 2,978 |
| 1151_Rep4 | 3,640 |
| mCherry_Rep1 | 287 |
| mCherry_Rep2 | 892 |
| mCherry_Rep3 | 878 |
| mCherry_Rep4 | 1,609 |
| **Total** | **11,527** |

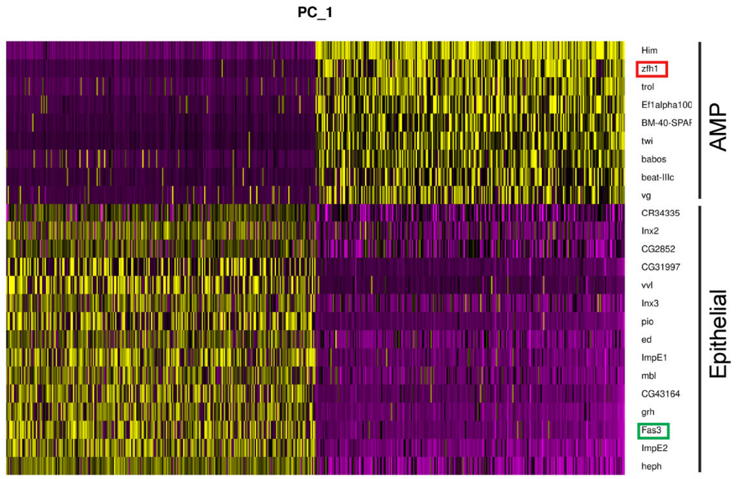


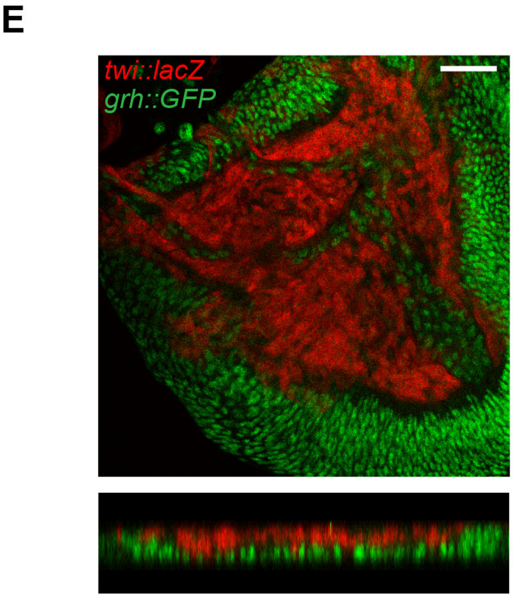


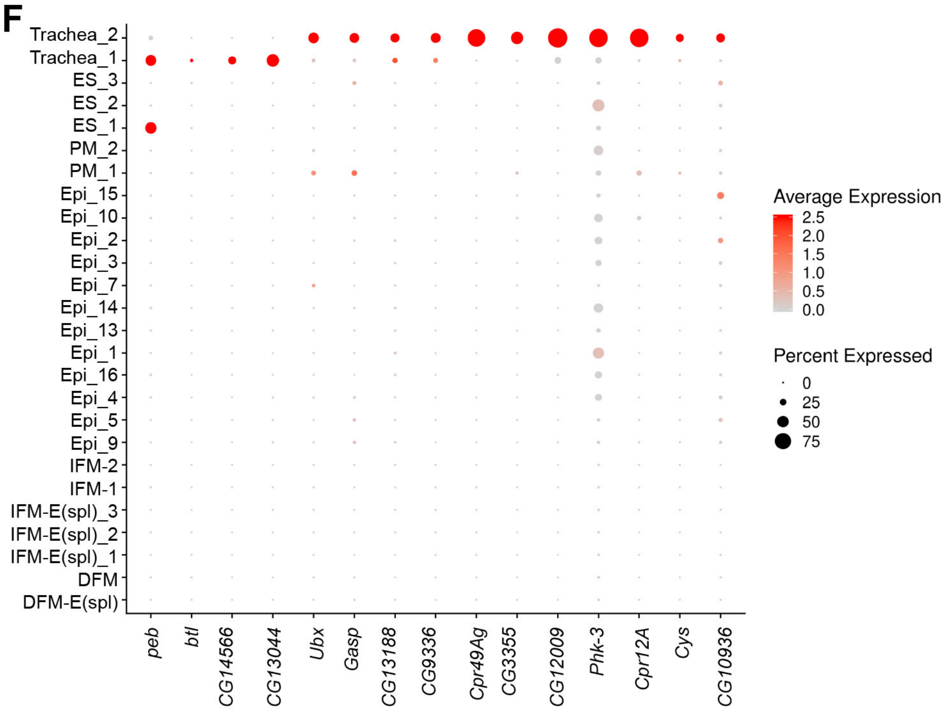

Sup Fig_S2

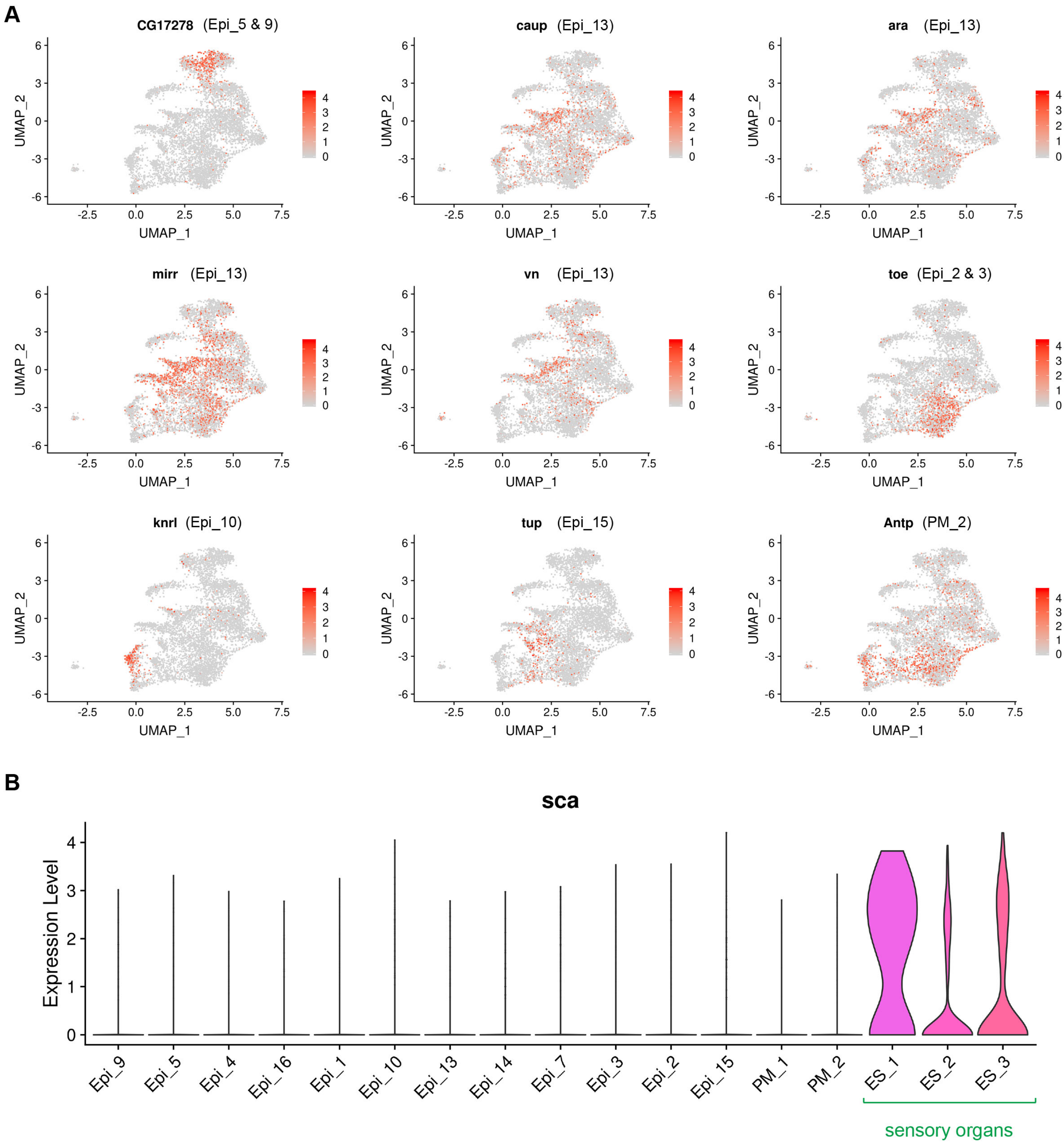

A
CG17278 (Epi_5 & 9)
caup (Epi_13)
ara (Epi_13)
mirr (Epi_13)
vn (Epi_13)
toe (Epi_2 & 3)
knrl (Epi_10)
tup (Epi_15)
Antp (PM_2)
UMAP_1
UMAP_2
B
sca
Expression Level
Epi_9
Epi_5
Epi_4
Epi_16
Epi_1
Epi_10
Epi_13
Epi_14
Epi_7
Epi_3
Epi_2
Epi_15
PM_1
PM_2
ES_1
ES_2
ES_3
sensory organs

Sup Figure S3

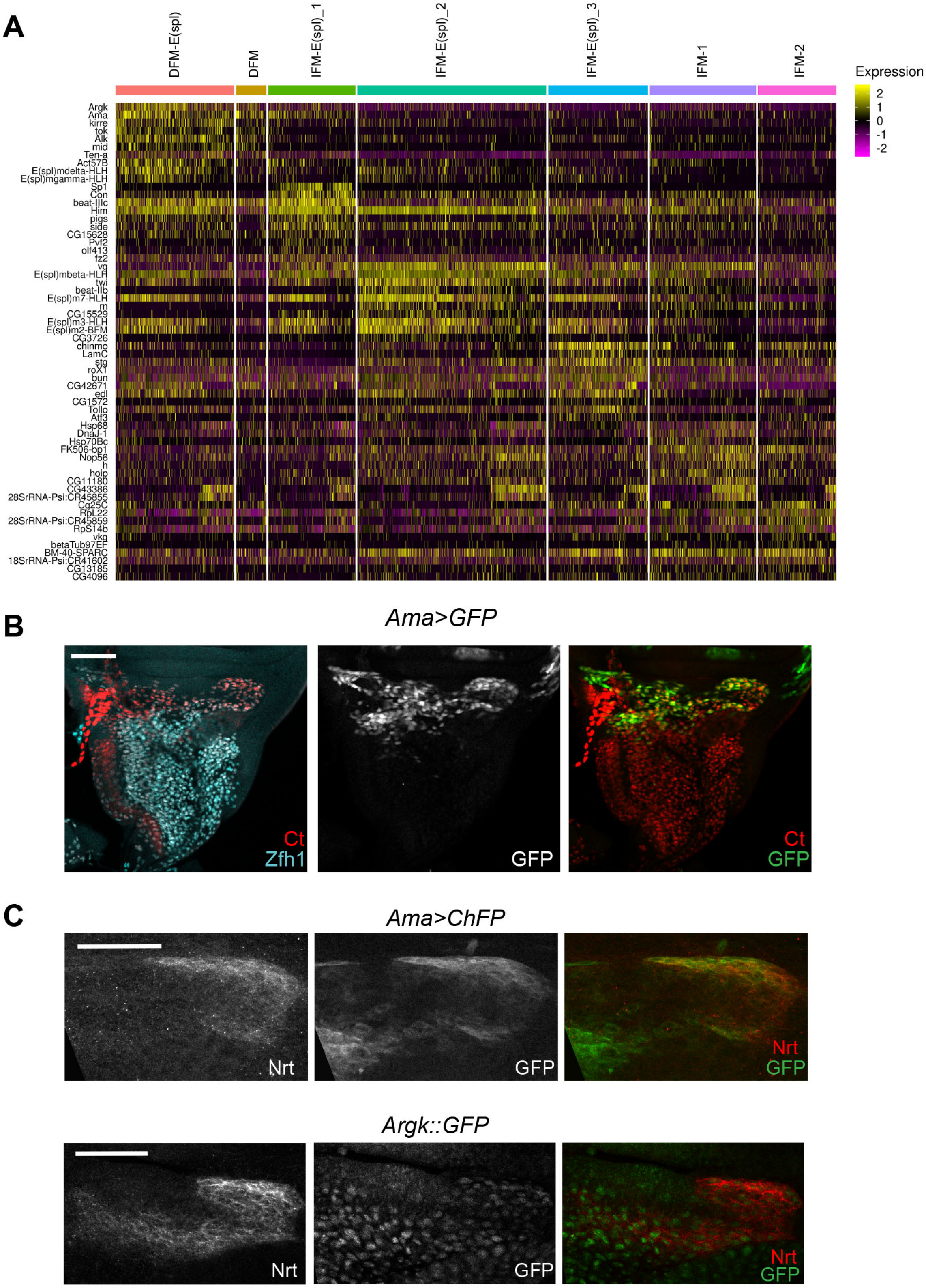

Sup Figure S4

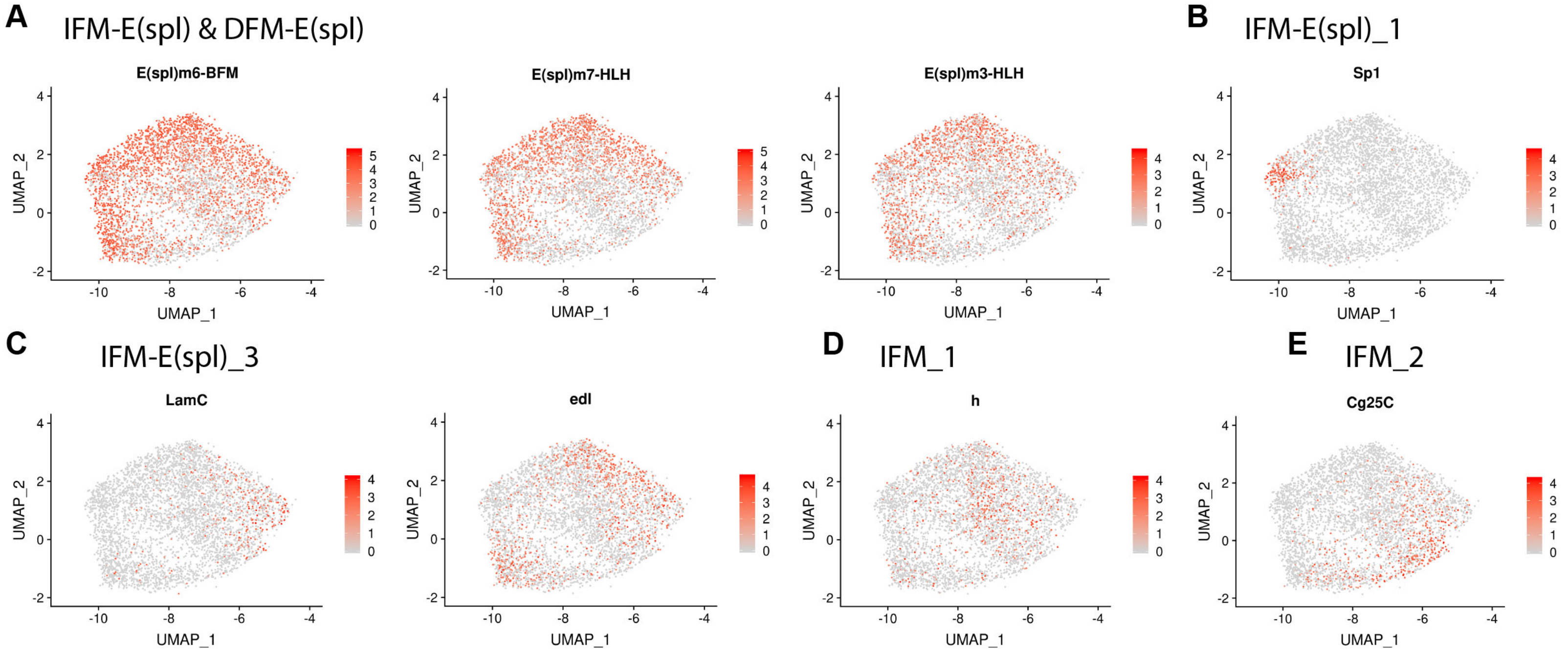

Sup Figure S5

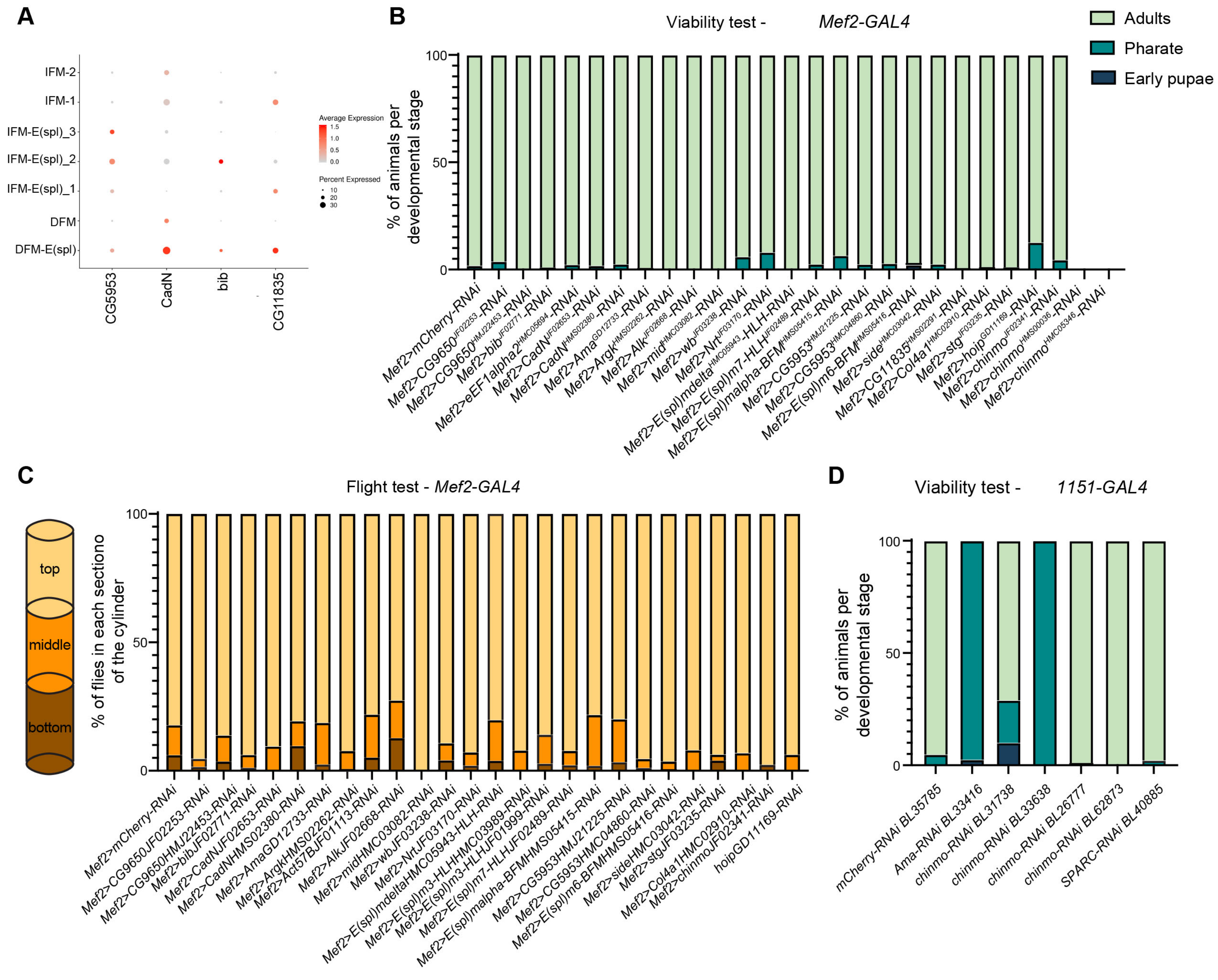

Sup Figure S6

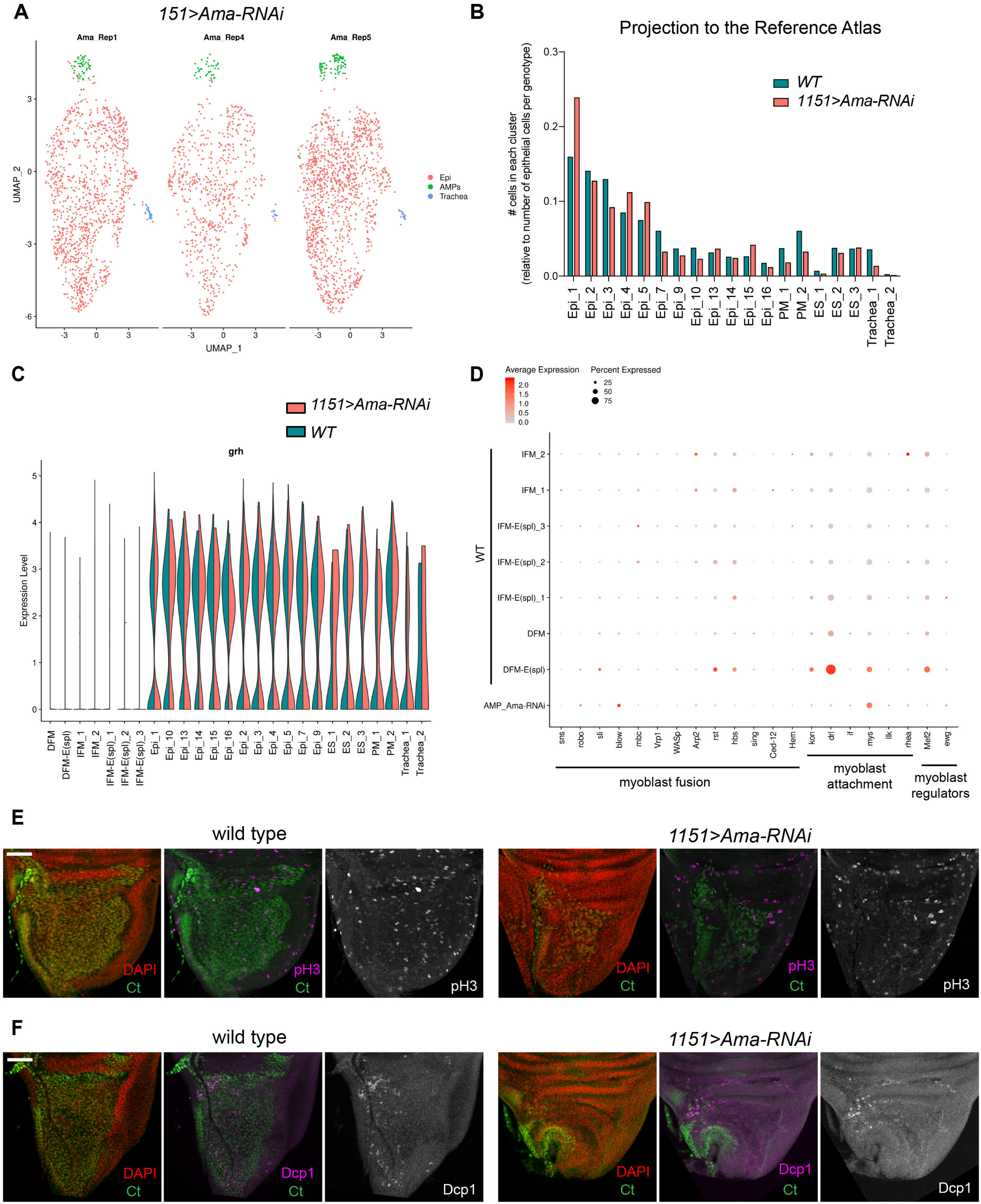

### Sup Figure S7

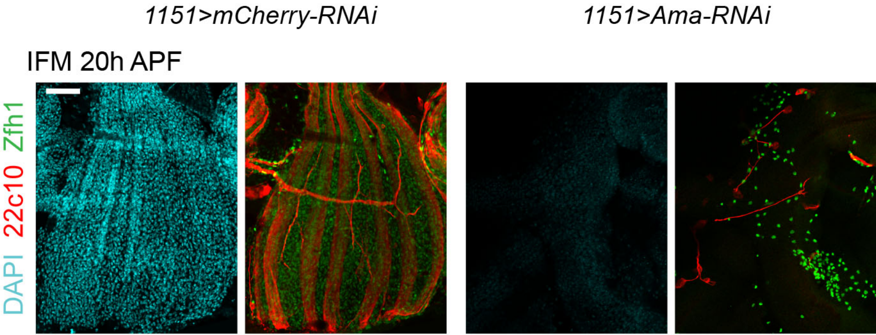
